# CAR-macrophage immunotherapy targeting fibroblast-like synoviocytes ameliorates inflammatory and degenerative arthritis

**DOI:** 10.64898/2026.10.25.761728

**Authors:** Yangli Xu, Zhuqian Wang, Duoli Xie, Luo Zhang, Xiushan Yin, Chao Liang

## Abstract

Rheumatoid arthritis (RA) is an inflammatory arthritis, whereas osteoarthritis (OA) is a degenerative joint disease. Despite their distinct etiologies, both are characterized by the pathogenic activation of fibroblast-like synoviocytes (FLS), which contribute to synovial hyperplasia, persistent inflammation, and cartilage destruction. Accordingly, pathogenic FLS represent an attractive therapeutic target for both RA and OA. Macrophages exhibit remarkable phenotypic plasticity, and interleukin-4 (IL-4) promotes an anti-inflammatory, tissue-reparative M2-like phenotype. Fibroblast activation protein α (FAP) and nucleolin (NCL) are highly expressed on the surface of pathological FLS while exhibiting limited accessibility in normal tissues. Here, we developed two IL-4-armored, FLS-targeted chimeric antigen receptor macrophage (CAR-M) platforms directed against FAP or NCL, termed CAR-M2-FAP and CAR-M2-NCL, respectively. *In vitro*, both platforms enhanced FLS phagocytosis and promoted an M2-like transcriptional profile. Transcriptomic analysis further revealed enrichment of phagocytic, anti-inflammatory, and tissue-reparative programs. In a mouse model of RA, both CAR-M2 platforms attenuated arthritis severity, synovial hyperplasia, cartilage loss, and bone erosion. Similarly, in a mouse model of OA, they reduced synovitis, cartilage degeneration, and osteophyte formation. No overt acute toxicity was observed in either RA or OA mice following treatment with CAR-M2-FAP or CAR-M2-NCL. Collectively, the concordant therapeutic effects of CAR-M2-FAP and CAR-M2-NCL across RA and OA models provide preclinical proof of concept for FLS-targeted CAR-M2 as a potential therapeutic strategy for inflammatory and degenerative arthritis.

**Graphical Abstract:** 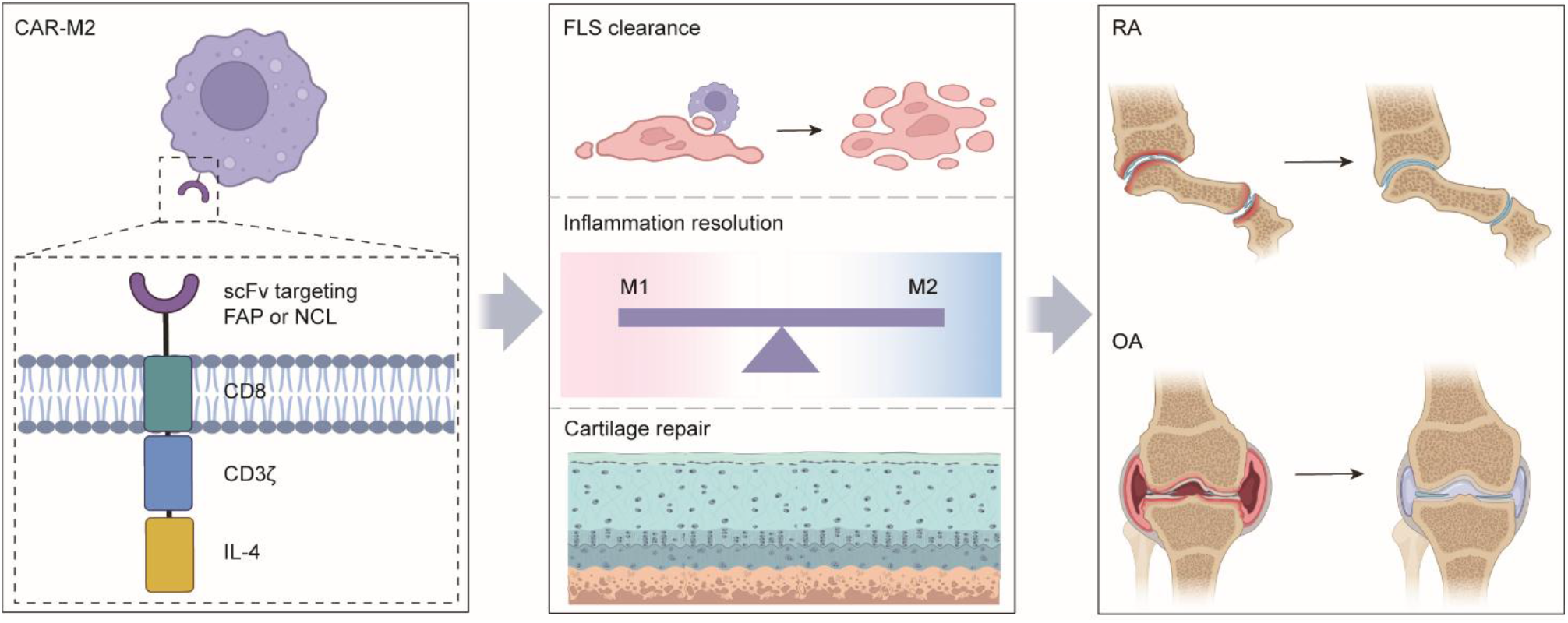

## Introduction

Arthritis is a common musculoskeletal disorder characterized by pain, inflammation and progressive joint dysfunction, and encompasses more than 100 distinct entities^1^. Among them, rheumatoid arthritis (RA) and osteoarthritis (OA) are the two most prevalent forms. RA is a chronic systemic autoimmune disease that primarily affects the small joints of the hands and feet, whereas OA is a heterogeneous whole-joint disorder that most commonly involves weight-bearing joints, particularly the knees and hips^2,3^. Despite their distinct etiologies, RA and OA share several pathological features, including immune cell infiltration, synovial hyperplasia and cartilage destruction^4,5^. Current treatments for RA, including disease-modifying antirheumatic drugs (DMARDs), have substantially improved disease management^6^. Despite this, a large proportion of RA patients still do not respond adequately to DMARDs^4,7^. In OA, disease-modifying osteoarthritis drugs (DMOADs) remain under active development, and no DMOADs have yet been approved for clinical use^8,9^. Thus, there is an urgent need for therapeutic strategies that address the unmet clinical demand for RA and OA treatment.

Fibroblast-like synoviocytes (FLS) are highly specialized mesenchymal cells residing in the intimal lining of the synovium^10–12^. Under physiological conditions, they maintain joint homeostasis by producing extracellular matrix components and key constituents of synovial fluid, thereby supporting cartilage lubrication and nutrition supply^13^. In RA and OA, however, FLS undergo pathological activation and acquire aggressive phenotypes characterized by aberrant proliferation, enhanced migratory and invasive capacities, and resistance to apoptosis, thereby sustaining synovial inflammation and contributing to disease progression^14–17^. Activated FLS further promote extracellular matrix degradation and pathological tissue remodeling^18,19^. In RA, these pathogenic activities drive pannus formation, cartilage erosion, and bone destruction, whereas in OA, they contribute to cartilage degeneration, subchondral bone remodeling, and osteophyte formation^20–22^. Given their central role in orchestrating inflammatory responses and tissue destruction across both RA and OA, pathogenic FLS have emerged as an attractive therapeutic target, and FLS-targeted strategies represent a promising approach for the treatment of inflammatory and degenerative arthritis^16,19,23^.

Chimeric antigen receptor (CAR)-engineered cellular therapies have expanded beyond oncology and are increasingly being investigated for the treatment of autoimmune and inflammatory diseases^24,25^. Macrophages are particularly attractive effector cells because their intrinsic phagocytic capacity enables efficient target cell engulfment and clearance^26,27^. Moreover, macrophages exhibit remarkable phenotypic plasticity and can adopt distinct functional states in response to environmental cues^28,29^. Classically activated M1 macrophages are generally associated with pro-inflammatory responses and tissue injury, whereas M2 macrophages exert anti-inflammatory effects, facilitate extracellular matrix remodeling and support tissue repair^30,31^. In both RA and OA, macrophages are skewed toward an M1-dominant state, resulting in disruption of macrophage homeostasis and contributing to persistent inflammation and joint pathology^32–34^. Interleukin-4 (IL-4) has demonstrated anti-inflammatory and tissue-reparative effects in experimental models of arthritis^35,36^. Consistently, targeted delivery of *Il4* to bone marrow-derived macrophages (BMDMs) promotes M2 polarization^37^, while incorporation of *Il4* into macrophage-based constructs further reinforce the M2-like phenotype^38^. Together, these findings provide a rationale for developing FLS-targeted, IL-4-armored CAR-M2 macrophages as a therapeutic strategy for arthritis^39^.

Fibroblast activation protein α (FAP) and nucleolin (NCL) are highly expressed on the surface of pathological FLS, while exhibiting limited accessibility in most normal tissues, making them attractive targets for FLS-directed therapy in RA and OA^17,40^. Here, we generated IL-4-armored, FAP-targeted CAR-M2 macrophages (CAR-M2-FAP) and NCL-targeted CAR-M2 macrophages (CAR-M2-NCL) by transducing BMDMs with CAR constructs incorporating an IL-4 expression module. Both CAR-M2 platforms mediated antigen-dependent phagocytosis of pathogenic FLS and suppressed inflammatory responses while exhibiting tissue-reparative ability *in vitro*. Following administration in RA and OA models, CAR-M2-FAP and CAR-M2-NCL preferentially accumulated in inflamed joints, interacted with pathogenic FLS, alleviated disease severity, and preserved cartilage and bone integrity. Importantly, blockade of the corresponding CAR antigens with recombinant proteins markedly abrogated these therapeutic effects, supporting the antigen-dependent nature of CAR-M2 activity. CAR-M2 treatment was well tolerated, with no overt systemic or organ toxicity detected in either disease model. Collectively, these findings establish the feasibility of FLS-targeted CAR-M2 therapy and highlight its potential as a therapeutic strategy for both inflammatory and degenerative arthritis.

## Results

### Workflow of FLS-targeted CAR-M2 construction and therapeutic evaluation in arthritis

To establish FLS-targeted CAR-M2 platforms for the treatment of arthritis, BMDMs were transduced with CAR constructs targeting FAP or NCL and incorporating IL-4 to promote an anti-inflammatory and tissue-reparative macrophage phenotype. The resulting FAP- and NCL-targeted CAR-M2 cells, designated CAR-M2-FAP and CAR-M2-NCL, respectively, were subsequently characterized and evaluated in mouse models of RA and OA (**Fig. 1A**). The proposed therapeutic mechanism is illustrated in **Fig. 1B**. Upon recognition of FAP- or NCL-expressing pathogenic FLS, CAR-M2 cells mediate antigen-dependent phagocytic clearance of target cells. Meanwhile, IL-4 signaling promotes an M2-like, anti-inflammatory and tissue-reparative state, thereby suppressing synovial inflammation and pathological tissue remodeling and preserving cartilage and bone integrity. Collectively, this approach combines antigen-specific FLS recognition and phagocytic clearance with IL-4-mediated macrophage reprogramming, providing a dual-action therapeutic strategy for both inflammatory and degenerative arthritis.

**Figure 1.**
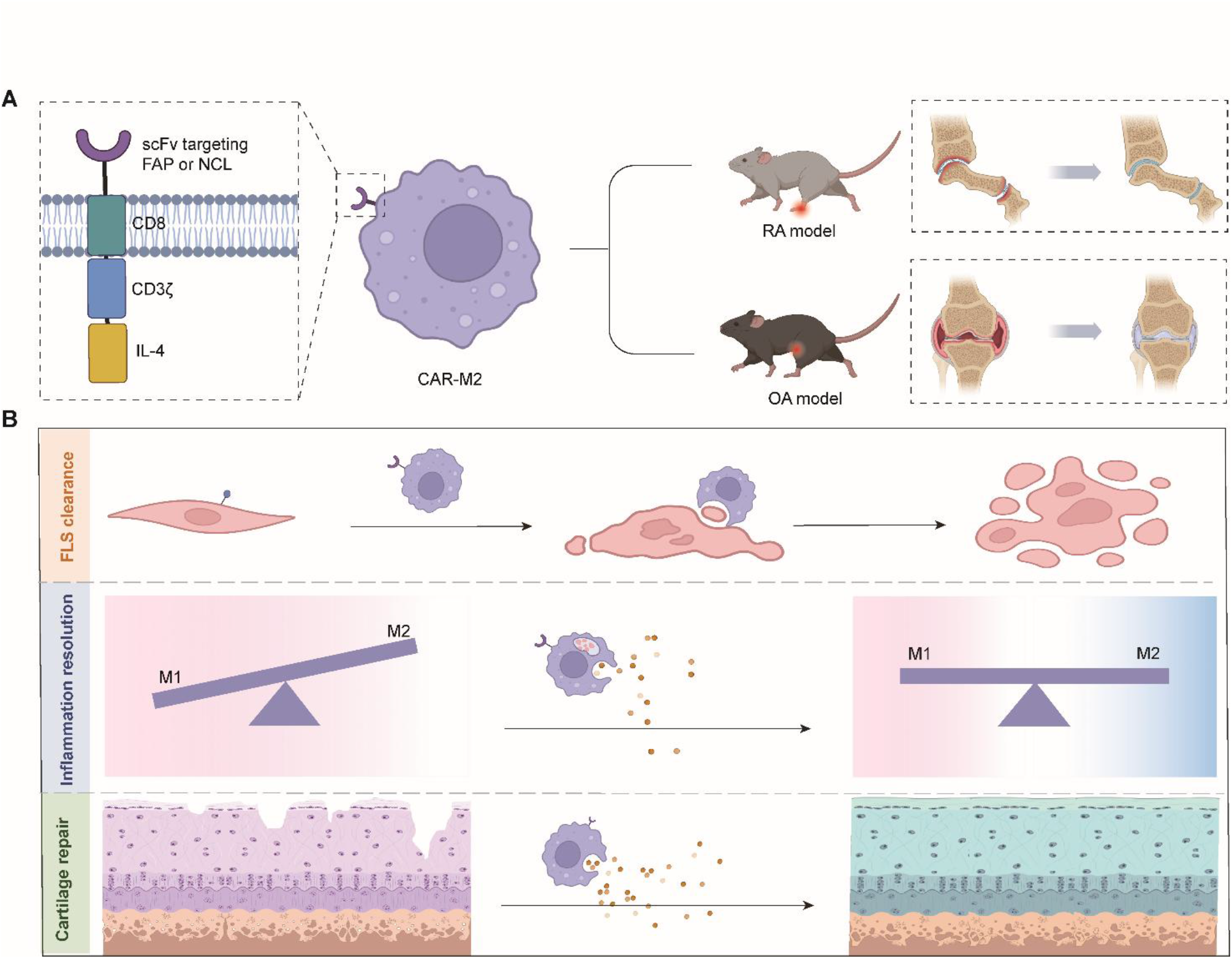
Construction and therapeutic evaluation of FLS-targeted CAR-M2 platforms in arthritis. **(A)** Schematic overview of CAR-M2-FAP and CAR-M2-NCL construction and therapeutic evaluation. BMDMs were engineered with CAR constructs targeting FAP or NCL and incorporating an IL-4 expression module to generate IL-4-armored CAR-M2-FAP and CAR-M2-NCL cells. The engineered macrophages were subsequently characterized and evaluated in mouse models of RA and OA. **(B)** Proposed mechanism of action of FLS-targeted CAR-M2 therapy. CAR-M2 cells recognize pathogenic FLS through CAR-mediated engagement of FAP or NCL and promote their antigen-dependent phagocytic clearance, while IL-4 signaling reinforces an anti-inflammatory and tissue-reparative macrophage phenotype. These coordinated activities are proposed to suppress synovial inflammation and pathological tissue remodeling, thereby preserving cartilage and bone integrity.

### Construction and *in vitro* characterization of CAR-M2-FAP

The CAR-M2-FAP construct comprised an N-terminal CD8 signal peptide (SP), a FAP-specific single-chain variable fragment (scFv), a CD8 transmembrane domain (TM), a CD3ζ intracellular signaling domain containing immunoreceptor tyrosine-based activation motifs (ITAMs) and a T2A-split IL-4-secreting domain. A Myc tag was incorporated for CAR detection, while green fluorescent protein (GFP) was co-expressed as a reporter to facilitate cell tracking and enrichment. BMDMs without transduction served as untransduced controls (BMDM-UTD), whereas BMDMs transduced with a CAR construct lacking both the scFv and IL-4-secreting domain served as the vector control (BMDM-vector) (**Fig. 2A**). Western blot analysis confirmed successful CAR expression in transduced BMDMs (**Fig. 2B**). Enzyme-linked immunosorbent assays (ELISA) demonstrated significantly increased IL-4 secretion by CAR-M2-FAP compared with BMDM-UTD or BMDM-vector (**Fig. 2C**). A Glycine₄-Serine (G4S) linker was included to ensure structural flexibility and proper folding of the CAR protein^41^. Flow cytometry further demonstrated robust CAR expression, with 95.1% of CAR-M2-FAP cells exhibiting a detectable G4S signal, markedly higher than that observed in BMDM-UTD or BMDM-vector (**Fig. 2D** and **2E**).

**Figure 2.**
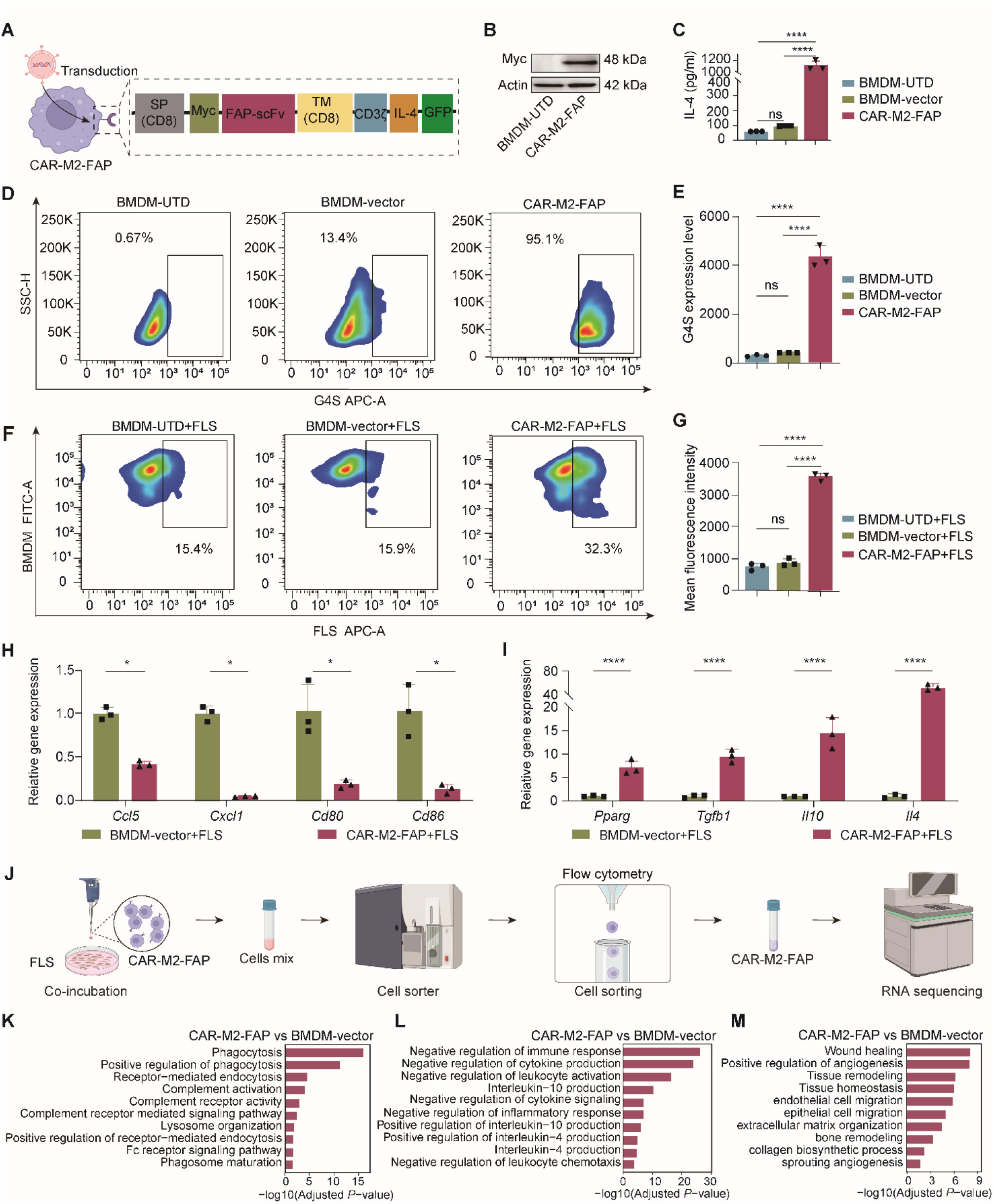
Generation and functional characterization of CAR-M2-FAP. **(A)** Schematic illustration of the lentiviral construct encoding CAR-M2-FAP. **(B)** Western blot analysis of Myc-tagged CAR expression in BMDM-UTD and CAR-M2-FAP. **(C)** ELISA analysis of IL-4 secretion in the culture supernatants of BMDMs following transduction. **(D** and **E)** Representative flow cytometry plots **(D)** and quantification **(E)** of G4S expression in BMDMs. **(F** and **G)** Representative flow cytometry plots **(F)** and quantification **(G)** of pathogenic FLS phagocytosis by BMDM-UTD, BMDM-vector, and CAR-M2-FAP following 12 h of co-culture at an effector-to-target (E:T) ratio of 3:1. **(H** and **I)** RT-qPCR analysis of M1-associated/pro-inflammatory genes **(H)** and M2-associated/anti-inflammatory genes **(I)** in BMDMs following co-culture with pathogenic FLS. **(J)** Schematic workflow of RNA-seq analysis of BMDM-vector and CAR-M2-FAP following co-culture with pathogenic FLS. **(K-M)** GO enrichment analysis of biological processes associated with phagocytosis **(K)**, inflammatory regulation **(L)**, and tissue repair **(M)** in CAR-M2-FAP versus BMDM-vector. Each experiment was independently repeated three times. Data are presented as mean ± SD. *P-*values were calculated by one-way ANOVA followed by Tukey’s post hoc test **(C, E,** and **G)** or paired Student’s t-test **(H** and **I)**: \**P* < 0.05, \*\*\*\**P* < 0.0001. ns, not significant.

We next evaluated the functional activity of CAR-M2-FAP by assessing its phagocytic capacity toward pathogenic mouse FLS. Following co-culture with FLS, flow cytometric analysis demonstrated significantly enhanced phagocytic activity in CAR-M2-FAP compared with both BMDM-UTD and BMDM-vector controls (**Fig. 2F** and **2G**). To determine whether CAR-M2-FAP acquired an anti-inflammatory and tissue-reparative transcriptional phenotype following interaction with pathogenic FLS, we performed reverse transcription quantitative PCR (RT-qPCR). Compared with BMDM-vector, CAR-M2-FAP exhibited significantly reduced expression of several pro-inflammatory M1-associated genes, including chemokine (C-C motif) ligand 5 (*Ccl5*), C-X-C motif chemokine ligand 1 (*Cxcl1*), *Cd80*, and *Cd86* (**Fig. 2H**). In contrast, genes associated with anti-inflammatory and tissue-reparative functions, including peroxisome proliferator-activated receptor gamma (*Pparg*), transforming growth factor beta 1 (*Tgfb1*), *Il10*, and *Il4*, were significantly upregulated (**Fig. 2I**). These findings indicate that CAR-M2-FAP not only enhances FLS phagocytosis but also exhibits a transcriptional profile consistent with an anti-inflammatory and tissue-reparative macrophage phenotype.

To further define the transcriptional programs underlying these functional changes, we performed bulk RNA sequencing (RNA-seq) of fluorescence-activated cell sorting (FACS)-purified CAR-M2-FAP cells following co-incubation with pathogenic FLS (**Fig. 2J**). Gene Ontology (GO) enrichment analysis revealed enhanced phagocytic programs in CAR-M2-FAP compared with BMDM-vector, with significant enrichment of processes associated with phagocytosis, receptor-mediated endocytosis, complement activation, lysosome organization, and phagosome maturation (**Fig. 2K**). In parallel, inflammatory programs were broadly attenuated, as indicated by enrichment of biological processes related to negative regulation of immune responses, cytokine production, leukocyte activation, cytokine signaling, and inflammatory responses, together with increased signatures associated with IL-4 and IL-10 production (**Fig. 2L**). Moreover, CAR-M2-FAP exhibited enrichment of tissue-reparative programs, including wound healing, angiogenesis, tissue remodeling, extracellular matrix organization, and collagen biosynthetic processes (**Fig. 2M**). Collectively, these transcriptomic findings demonstrate that CAR-M2-FAP integrates enhanced phagocytic activity with anti-inflammatory and tissue-reparative transcriptional programs following interaction with pathogenic FLS.

### Therapeutic potential of CAR-M2-FAP in CIA mice

Collagen-induced arthritis (CIA) is a widely used animal model of RA that recapitulates key immunological and pathological features of human disease^42^. To establish CIA, mice were immunized with bovine type II collagen (CII) emulsion in complete Freund’s adjuvant (CFA) on day 0. Mice with established CIA were subsequently treated with a single intravenous injection of phosphate-buffered saline (PBS), BMDM-UTD, BMDM-vector, CAR-M2-FAP, or CAR-M2-FAP pre-incubated with recombinant FAP (CAR-M2-FAP + rFAP) on day 28 (**Fig. 3A**). Following treatment, we performed arthritic scoring, micro-computerized tomography (μCT) scans, histopathological analyses, and immunofluorescence (IF) staining. CIA mice treated with CAR-M2-FAP showed markedly reduced arthritis scores compared with mice receiving PBS, BMDM-UTD, or BMDM-vector (**Fig. 3B**). IF staining revealed substantial co-localization of FAP and GFP signals in the joints of CAR-M2-FAP-treated mice, whereas pre-incubation with rFAP markedly reduced this co-localization (**Fig. 3C**), supporting FAP-dependent CAR engagement *in vivo*. Consistent with the improved clinical scores, μCT analysis revealed marked attenuation of bone erosion and joint destruction following CAR-M2-FAP treatment (**Fig. 3D**). Histopathological analysis showed that CAR-M2-FAP substantially attenuated synovial hyperplasia and cartilage loss compared with PBS-, BMDM-UTD-, and BMDM-vector-treated mice (**Fig. 3E** and **3F**). IF staining demonstrated reduced expression of CD80 and proportion of FAP-positive pathogenic FLS, together with enhanced expression of IL-4 and IL-10 in the joints of CAR-M2-FAP-treated mice (**Fig. 3G-3K**). Notably, pre-incubation with rFAP largely abrogated the therapeutic effects of CAR-M2-FAP, as evidenced by the reversal of improvements in arthritis severity, bone erosion, synovial hyperplasia, cartilage loss, and macrophage-associated markers (**Fig. 3B-3K**). These findings support the FAP-dependent therapeutic activity of CAR-M2-FAP *in vivo*. We next evaluated the *in vivo* safety profile of CAR-M2-FAP. Body weight remained stable throughout the treatment period in CAR-M2-FAP-treated mice **(Fig. S1A)**. Serum biochemical analysis revealed no significant abnormalities in parameters associated with hepatic function, including albumin (ALB), alkaline phosphatase (ALP), alanine aminotransferase (ALT), and aspartate aminotransferase (AST), or in renal function, including creatinine (Cr) and urea **(Fig. S1B-S1G)**. Hematoxylin and eosin (H&E) staining further revealed no overt histopathological abnormalities in the heart, liver, spleen, lung, or kidney **(Fig. S1H)**. Collectively, these findings demonstrate that systemic administration of CAR-M2-FAP alleviated CIA-associated joint pathology while exhibiting no detectable systemic or major-organ toxicity under the conditions tested.

**Figure 3.**
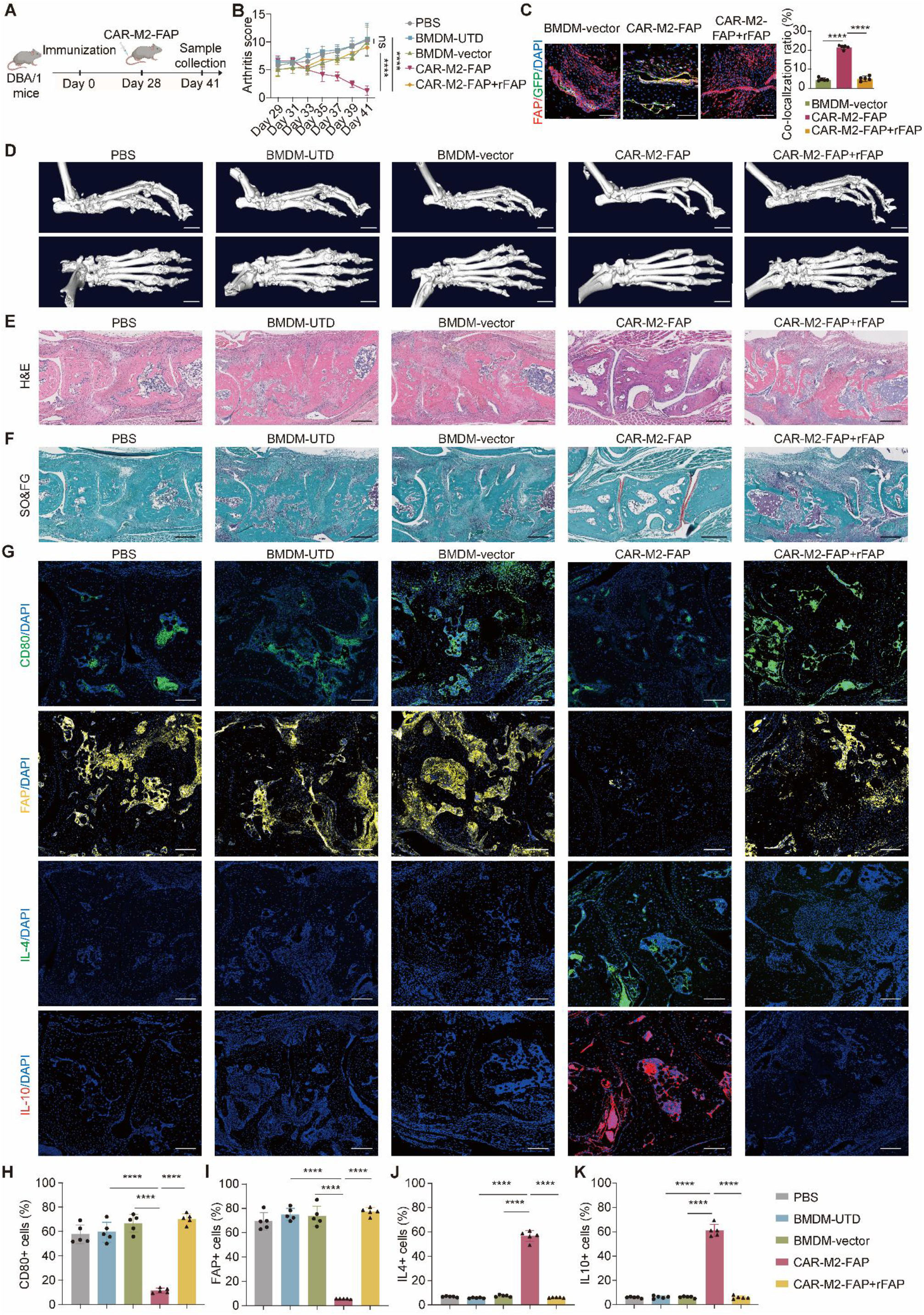
Therapeutic effects of CAR-M2-FAP in CIA mice. **(A)** Schematic illustration of CIA induction, CAR-M2-FAP treatment, and sample collection. **(B)** Clinical arthritis scores of CIA mice following intravenous administration of PBS, BMDM-UTD, BMDM-vector, CAR-M2-FAP, or CAR-M2-FAP + rFAP. **(C)** Representative IF images of paw sections showing FAP and GFP co-localization and its quantification analysis. Nuclei were counterstained with DAPI. Scale bar = 50 μm. **(D)** Representative three-dimensional μCT reconstructions of hind paws from the indicated treatment groups. Scale bar = 1 mm. **(E)** Representative H&E-stained paw sections from the indicated treatment groups. Scale bar, 200 μm. **(F)** Representative SO&FG-stained paw sections from the indicated treatment groups. Scale bar, 200 μm. **(G)** Representative IF staining of paw sections for CD80, FAP, IL-4, and IL-10. Scale bar = 50 μm. **(H-K)** Quantification of CD80-positive cells **(H)**, FAP-positive cells **(I)**, IL-4-positive cells **(J),** and IL-10-positive cells **(K)** in paw sections from different treatment groups. n = 5 per group. Data are presented as mean ± SD. *P*-values were calculated by two-way repeated-measures ANOVA followed by Tukey’s post hoc test **(B)** or one-way ANOVA followed by Tukey’s post hoc test **(C** and **H-K)**: \*\*\*\**P* < 0.0001, ns, not significant.

### CAR-M2-NCL generation and therapeutic evaluation in CIA mice

To determine whether the therapeutic concept could be extended to an alternative FLS-associated target, we next applied the CAR-M2 framework to NCL, another surface-exposed target on pathogenic FLS (**Fig. 4A**). Successful CAR expression in CAR-M2-NCL was verified by western blotting (**Fig. 4B**), and ELISA confirmed enhanced IL-4 secretion relative to BMDM-UTD and BMDM-vector controls (**Fig. 4C**). Flow cytometric analysis showed 64.4% of CAR-M2-NCL cells were positive for the G4S-associated signal, whereas substantially fewer cells were positive in the BMDM-UTD and BMDM-vector groups **(Fig. S2A** and **S2B)**. Following co-culture with pathogenic FLS, CAR-M2-NCL exhibited robust phagocytic activity, with 72.9% of cells being phagocytosis-positive, significantly higher than that observed in both control groups (**Fig. 4D** and **4E**). RT-qPCR analysis revealed a coordinated shift in gene expression toward an anti-inflammatory phenotype, characterized by reduced expression of pro-inflammatory M1-associated genes and increased expression of anti-inflammatory and M2-associated genes following exposure to pathogenic FLS **(Fig. S2C** and **S2D)**.

**Figure 4.**
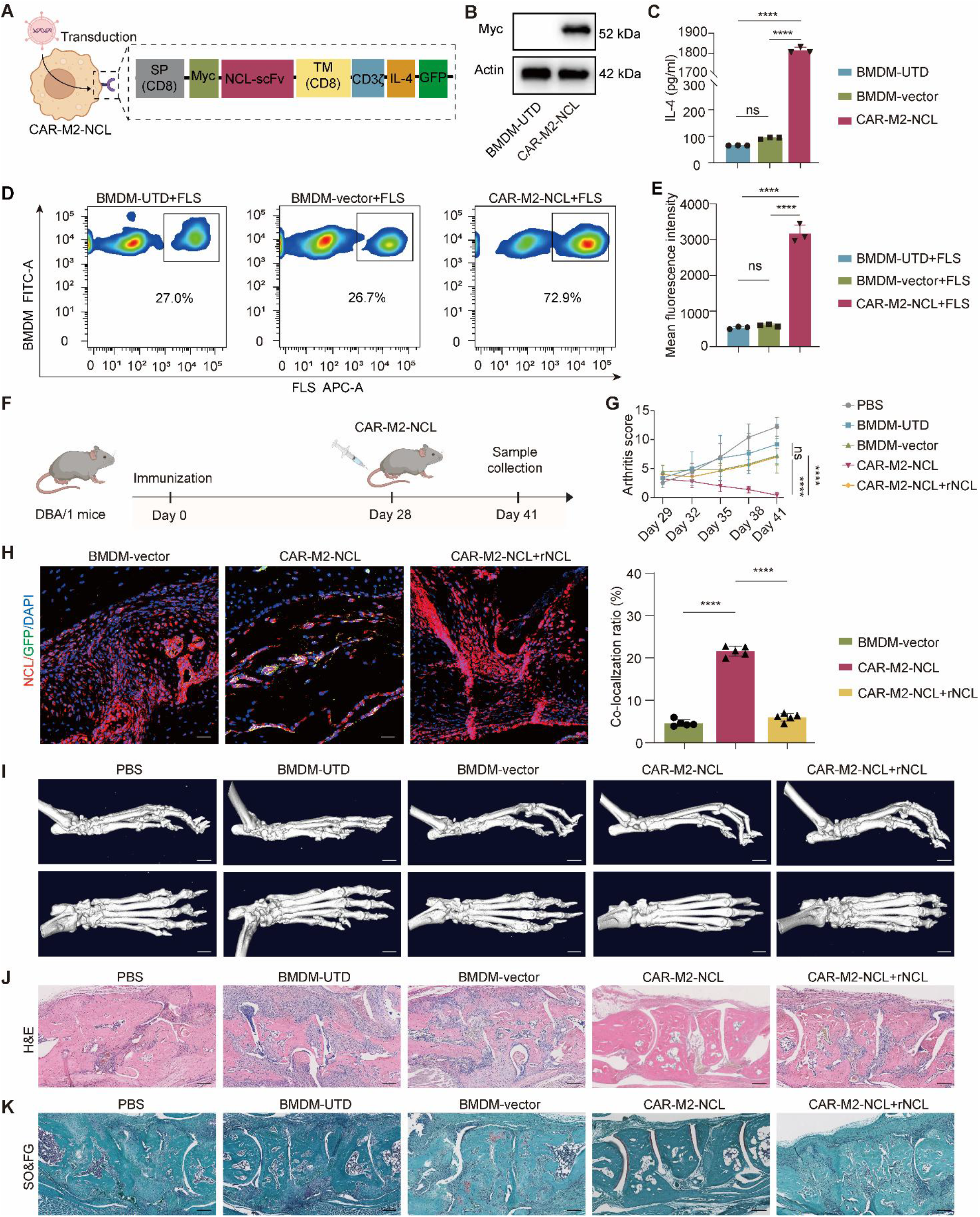
Generation and functional evaluation of CAR-M2-NCL. **(A)** Schematic illustration of the lentiviral construct encoding CAR-M2-NCL. **(B)** Western blot analysis of Myc-tagged CAR expression in BMDM-UTD and CAR-M2-NCL. **(C)** ELISA analysis of IL-4 secretion in the culture supernatants of BMDMs after transduction. **(D and E)** Representative flow cytometry plots **(D)** and quantification **(E)** of pathogenic FLS phagocytosis by BMDM-UTD, BMDM-vector, and CAR-M2-NCL following 12 h of co-culture at an E: T ratio of 3:1. **(F)** Schematic illustration of CIA induction, CAR-M2-NCL administration, and sample collection. **(G)** Clinical arthritis scores of CIA mice after intravenous injection of PBS, BMDM-UTD, BMDM-vector, CAR-M2-NCL, or CAR-M2-NCL + rNCL. **(H)** Representative IF images of paw sections showing NCL and GFP co-localization, with quantification of the co-localization ratio. Nuclei were counterstained with DAPI. Scale bar = 50 μm. **(I)** Representative three-dimensional μCT reconstructions of hind paws from different treatment groups. Scale bar = 1 mm. **(J** and **K)** H&E **(J)** and SO&FG staining **(K)** of paw sections from different treatment groups. Scale bar = 200 μm. *In vitro* experiments were independently repeated three times. For *in vivo* experiments, n = 5 mice per group. Data are presented as mean ± SD. *P-*values were calculated by one-way ANOVA followed by Tukey’s post hoc test **(C**, **E,** and **H)** or two-way repeated-measures ANOVA followed by Tukey’s post hoc test **(G)**: \**P* < 0.05, \*\**P* < 0.01, \*\*\**P* < 0.001, \*\*\*\**P* < 0.0001, ns, not significant.

We next evaluated the therapeutic efficacy of CAR-M2-NCL *in vivo*. Mice with established CIA received a single intravenous injection of the indicated BMDM preparations on day 28 (**Fig. 4F**). CAR-M2-NCL treatment resulted in a significant reduction in clinical arthritis scores relative to PBS-, BMDM-UTD-, and BMDM-vector-treated mice (**Fig. 4G**). IF analysis showed increased overlap between NCL and GFP signals in the joints of CAR-M2-NCL-treated mice, whereas pre-incubation with recombinant NCL (rNCL) markedly diminished this co-localization (**Fig. 4H**). Structural assessment by μCT revealed less extensive bone erosion and joint destruction following CAR-M2-NCL treatment (**Fig. 4I**). Histopathological analysis confirmed reduced synovial hyperplasia and cartilage loss in CAR-M2-NCL-treated mice (**Fig. 4J and 4K**). At the molecular level, CAR-M2-NCL treatment was associated with reduced CD80 expression and a decreased proportion of NCL-positive pathogenic FLS, together with elevated levels of IL-4 and IL-10 in joint tissues **(Fig. S3A-S3E)**. The target dependence of these therapeutic effects was further supported by rNCL competition. Pre-incubation of CAR-M2-NCL with rNCL significantly weakened its effects on clinical arthritis, bone erosion, synovial hyperplasia, cartilage preservation, and macrophage-associated markers (**Fig. 4G-4K and Fig. S3A-S3E)**. CAR-M2-NCL-treated mice maintained stable body weights throughout the observation period **(Fig. S4A)**, while serum biochemical analyses revealed no apparent abnormalities indicative of hepatic or renal injury **(Fig. S4B-S4G)**. Moreover, H&E staining showed no overt histopathological abnormalities in major organs **(Fig. S4H)**. Collectively, these findings demonstrate that CAR-M2-NCL efficiently mediates antigen-dependent phagocytosis of pathogenic FLS *in vitro* and exerts target-dependent therapeutic effects in CIA mice, while exhibiting a favorable safety profile.

### Therapeutic potential of CAR-M2-FAP and CAR-M2-NCL in OA mice

We next determined whether the CAR-M2 platform could be applied to OA. Destabilization of the medial meniscus (DMM) is a commonly used mouse model that recapitulates key pathological features of OA^43^. Following DMM surgery on day 0, mice received right-knee intra-articular injection of BMDM-UTD, BMDM-vector, CAR-M2-FAP, CAR-M2-FAP + rFAP, CAR-M2-NCL, or CAR-M2-NCL + rNCL on day 14 (**Fig. 5A**). We examined the joint localization of the engineered macrophages. IF staining showed substantial co-localization of GFP with FAP or NCL in the corresponding CAR-M2-treated joints, whereas pre-incubation with rFAP or rNCL markedly reduced the respective FAP-GFP or NCL-GFP co-localization ratios (**Fig. 5B-5D**). μCT analysis of the three-dimensional reconstruction of the knee joints showed that both CAR-M2-FAP and CAR-M2-NCL markedly reduced joint deformity and osteophyte formation compared with the control groups (**Fig. 5E**). Histopathological examination further revealed attenuated synovitis and cartilage loss following treatment with either CAR-M2-FAP or CAR-M2-NCL (**Fig. 5F** and **5G**). These improvements were consistently reflected by quantitative measurements, including the Osteoarthritis Research Society International (OARSI) score, cartilage area, osteophyte score, and synovitis score, all of which indicated reduced OA severity in CAR-M2-treated mice (**Fig. 5H-5K**). IF staining demonstrated increased COL2A1, IL-4, and IL-10 signals, accompanied by reduced CD80 expression and proportions of FAP-positive and NCL-positive pathogenic FLS, in the joints of CAR-M2-treated mice (**Fig. 6A-6G**). However, the structural, histopathological, and quantitative benefits of CAR-M2 treatment were substantially attenuated by pre-incubation with rFAP or rNCL (**Fig. 5E-5K and Fig. 6A-6G**), supporting the target-dependent activity of both CAR-M2 platforms. Body weight remained stable throughout the treatment period in mice receiving CAR-M2-FAP or CAR-M2-NCL **(Fig. S5A)**. Serum biochemical analyses revealed no apparent treatment-related hepatic and renal abnormalities **(Fig. S5B-S5G)**. Consistently, H&E staining of the heart, liver, spleen, lung, and kidney showed no histopathological abnormalities **(Fig. S5H)**. Collectively, these findings demonstrate that both FAP- and NCL-targeted CAR-M2 platforms attenuate structural, histopathological, and inflammatory features of OA in DMM mice while exhibiting no detectable systemic or major-organ toxicity under the conditions tested.

**Figure 5.**
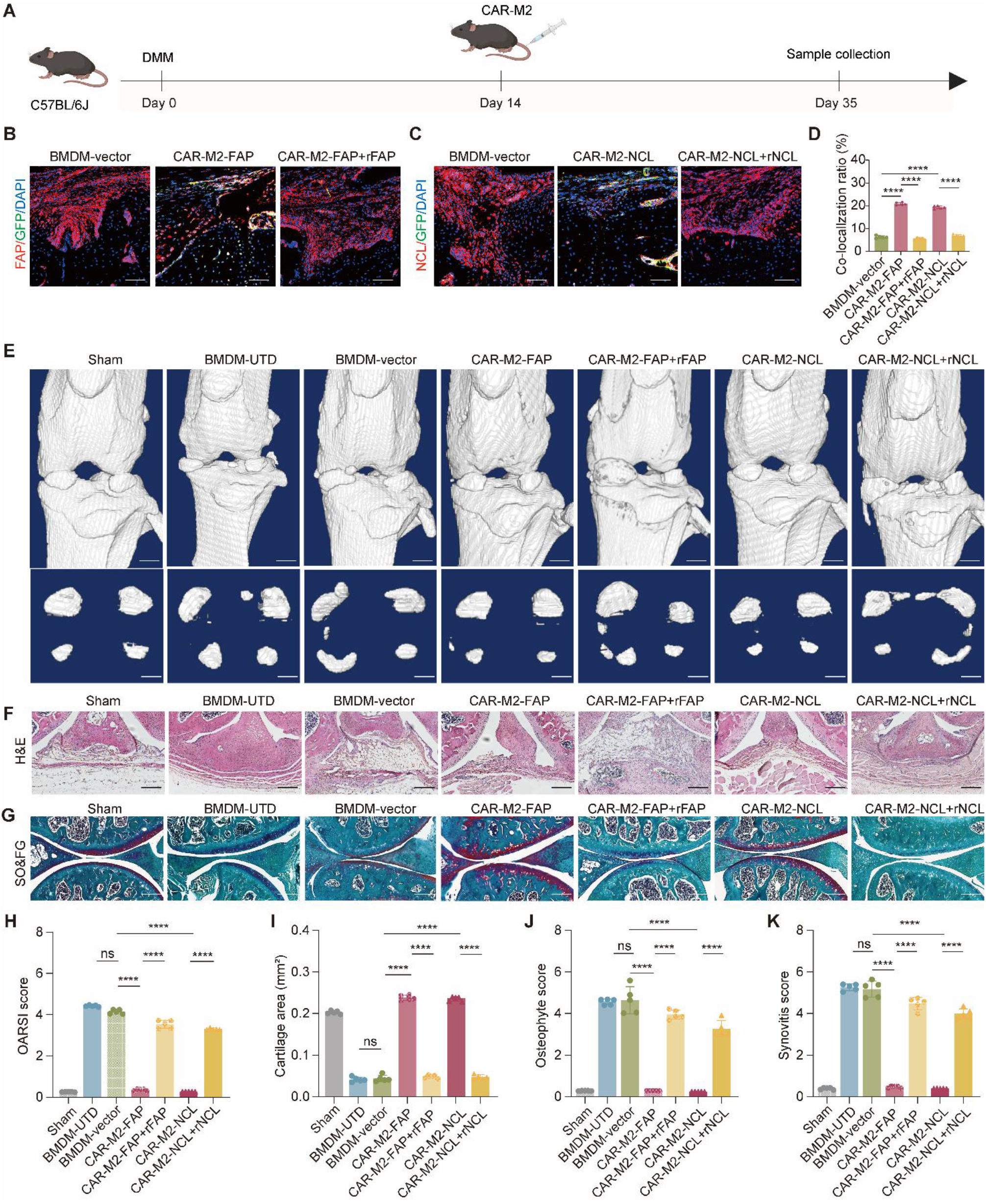
Therapeutic effects of CAR-M2 in DMM mice. **(A)** Schematic diagram of DMM surgery, CAR-M2 administration, and sample collection. **(B-D)** Representative IF staining of knee sections showing FAP/GFP **(B)** or NCL/GFP **(C)** co-localization, with quantification of the co-localization ratio **(D)**. Scale bar = 50 μm. **(E)** Representative three-dimensional μCT reconstructions of knee joints and osteophytes from sham or DMM mice after different treatments. Scale bar = 300 μm. **(F)** Representative H&E staining of knee sections from different treatment groups. Scale bar = 50 μm. **(G)** Representative SO&FG staining of knee sections from different treatment groups. Scale bar = 200 μm. **(H-K)** Quantification of OARSI score **(H)**, cartilage area **(I)**, osteophyte score **(J)**, and synovitis score **(K)**. Data are presented as mean ± SD. n = 5 per group. *P-*values were calculated by one-way ANOVA followed by Tukey’s post hoc test **(D** and **H-K)**: \*\*\*\**P* < 0.0001, ns, not significant.

**Figure 6.**
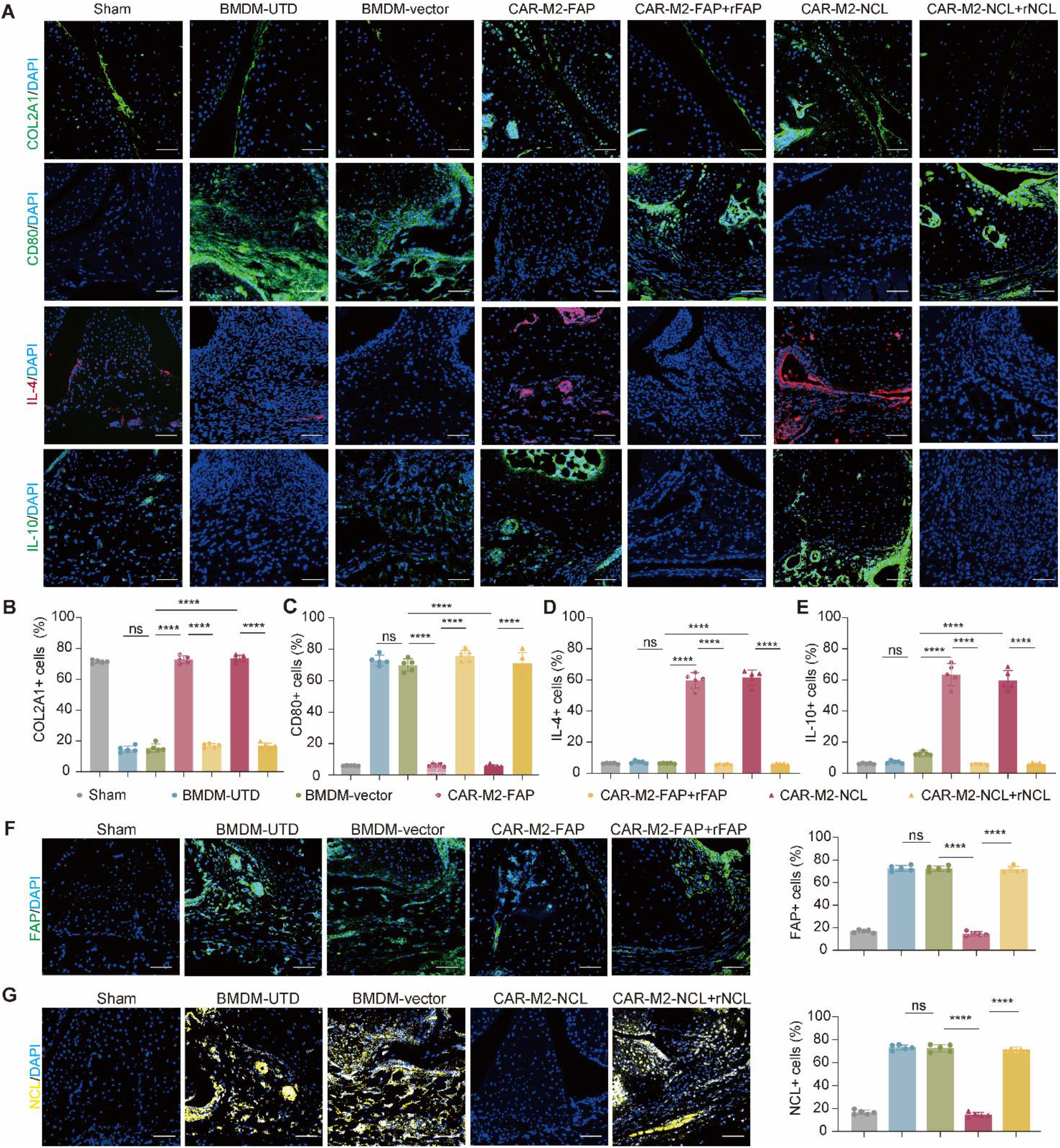
IF analysis of CAR-M2-treated DMM mice. **(A)** Representative IF staining of knee sections from sham or DMM mice after different treatments using antibodies against COL2A1, CD80, IL-4, and IL-10. Scale bar = 50 μm. **(B-E)** Quantification of COL2A1-positive cells **(B)**, CD80-positive cells **(C)**, IL-4-positive cells **(D),** and IL-10-positive cells **(E)** in knee sections from different treatment groups. **(F)** Representative IF staining and quantification of FAP-positive cells in knee sections from different treatment groups. Scale bar = 50 μm. **(G)** Representative IF staining and quantification of NCL-positive cells in knee sections from different treatment groups. Scale bar = 50 μm. Nuclei were counterstained with DAPI. Data are presented as mean ± SD. n = 5 per group. *P*-values were calculated by one-way ANOVA followed by Tukey’s post hoc test **(B-G)**: \*\*\*\**P* < 0.0001, ns, not significant.

## Discussion

CAR-M has emerged as a promising myeloid cell-based therapeutic platform that integrates programmable antigen recognition with the intrinsic phagocytic and immunomodulatory functions of macrophages. Although most CAR-M studies to date have focused on solid tumors and, more recently, fibrotic diseases^26,38,44–53^, their potential in arthritis has remained largely unexplored. This is particularly relevant because pathogenic FLS are not merely bystanders in arthritis but active drivers of synovial inflammation, extracellular matrix degradation, and progressive joint destruction. In the present study, we developed two IL-4-armored CAR-M2 platforms targeting FAP or NCL and demonstrated their therapeutic activity across both inflammatory and degenerative arthritis models. *In vitro*, CAR-M2-FAP and CAR-M2-NCL enhanced antigen-dependent phagocytosis of pathogenic FLS and promoted an anti-inflammatory and tissue-reparative transcriptional state, with CAR-M2-FAP further showing enrichment of gene programs associated with phagocytosis, inflammatory regulation, and tissue repair. *In vivo*, both CAR-M2 platforms accumulated at pathogenic FLS-associated sites in arthritic joints and alleviated clinical, structural, and histopathological features of disease in CIA and DMM models. Importantly, pre-incubation with recombinant FAP or NCL reduced CAR-M2 localization and substantially attenuated the therapeutic effects, supporting the target-dependent nature of these responses. Neither platform produced overt systemic or major-organ toxicity under the conditions tested. Collectively, these findings establish a proof of concept for targeting pathogenic FLS with IL-4-armored CAR-M2 cells and suggest that this approach may simultaneously address inflammatory and tissue-destructive processes in arthritis.

A key feature of our strategy is the integration of targeted pathogenic-cell clearance with local immunoregulation within the same therapeutic cell. IL-4 is a well-established regulator of macrophage phenotype that promotes alternative activation and supports anti-inflammatory and tissue-reparative responses^54,55^. However, systemic administration of recombinant IL-4 is constrained by its limited persistence and the difficulty of achieving sustained, therapeutically relevant concentrations within diseased tissues^56^. By incorporating an IL-4 expression module into CAR-M cells, we sought to couple antigen-directed engagement with sustained local delivery of an immunoregulatory signal. Consistent with this design, both CAR-M2-FAP and CAR-M2-NCL exhibited reduced expression of pro-inflammatory, M1-associated genes and increased expression of anti-inflammatory and M2-associated genes following interaction with pathogenic FLS. Transcriptomic analysis further demonstrated that CAR-M2-FAP acquired coordinated programs associated with enhanced phagocytosis, attenuation of inflammatory responses, and tissue repair. These findings suggest that IL-4 armoring may reinforce the functional state of CAR-M cells while their CAR-mediated engagement provides spatially restricted interaction with pathogenic FLS. Thus, rather than relying solely on depletion of pathogenic FLS, our platform combines cellular clearance with local macrophage reprogramming, potentially providing complementary control over both the cellular and inflammatory components of joint pathology. This concept is consistent with a previous study of CAR-M therapy in renal fibrosis, in which engineered M2 macrophages engulfed FAP-positive fibroblasts, reduced fibrosis, and promoted revascularization^38^. Together, these observations support the broader utility of combining CAR-mediated target recognition with macrophage-based immunoregulatory functions. Future studies could further explore alternative or combinatorial payloads, such as IL-10, to determine whether distinct immunoregulatory programs can further improve therapeutic efficacy or durability^57^.

The selection of FAP and NCL provides an additional strength of the present study. FAP is a type II transmembrane serine protease with dipeptidyl peptidase and endopeptidase activities^58,59^, while NCL is a multifunctional RNA-binding phosphoprotein primarily localized in the nucleolus that regulates ribosome biogenesis, chromatin remodeling, and transcription^60^. Despite their distinct biological functions, both proteins can become prominently exposed on the surface of pathological FLS, while showing limited accessibility in most normal tissues^17,61,62^. This disease-associated surface accessibility provides an important rationale for their exploitation as therapeutic targets. Previous studies have shown that selective depletion of FAP-positive FLS can reduce synovial inflammation and protect against cartilage and bone damage in experimental arthritis^63^. In our previous work, we likewise developed an NCL-targeting cytotoxic aptamer that selectively recognized and eliminated pathogenic FLS^40^. These findings provided independent support for FAP and NCL as candidate FLS-associated targets and motivated us to determine whether the CAR-M2 platform could be adapted to both antigens. As expected, our results demonstrate that CAR-M2-FAP and CAR-M2-NCL both exhibited efficient phagocytosis of pathogenic FLS *in vitro* and accumulated at FAP- or NCL-positive sites within arthritic joints *in vivo*. Moreover, blockade with recombinant FAP or NCL reduced target-associated CAR-M2 localization and substantially weakened therapeutic efficacy, supporting a critical contribution of antigen-dependent CAR engagement to the observed activity. Importantly, the similar therapeutic profiles obtained with two mechanistically distinct FLS-associated targets argue against the effects being restricted to a single antigen and instead support the adaptability of the CAR-M2 framework to different pathogenic FLS surface markers. This target modularity may be particularly valuable in arthritis, where FLS are heterogeneous and may exhibit disease-, tissue-, and patient-specific differences in antigen expression.

The therapeutic implications of targeting pathogenic FLS extend beyond suppression of inflammation alone. Although RA and OA differ substantially in their initiating mechanisms and dominant pathological features, both diseases exhibit aberrant FLS activation, synovial remodeling, extracellular matrix disruption, and progressive joint damage^64^. In RA, pathogenic FLS contribute to pannus formation, inflammatory amplification, cartilage destruction, and bone erosion, whereas in OA, FLS participate in synovial inflammation, cartilage degeneration, subchondral bone remodeling, and osteophyte formation^65^. Our findings that both FAP- and NCL-targeted CAR-M2 cells reduced synovial inflammation and pathological structural changes in both CIA and DMM models therefore suggest that pathogenic FLS may represent a shared therapeutic node across inflammatory and degenerative arthritis. Notably, the effects observed in DMM mice were not limited to inflammatory markers but also included preservation of cartilage-associated COL2A1 and reductions in osteophyte formation, synovitis, and cartilage loss. These findings support the possibility that eliminating or functionally suppressing pathogenic FLS, while simultaneously promoting an anti-inflammatory and tissue-reparative macrophage state, may provide broader disease-modifying effects than approaches directed solely at inflammatory cytokines.

Several limitations should nevertheless be considered. First, although the efficacy of FAP- and NCL-targeted CAR-M2 cells was demonstrated in representative models of RA and OA, whether this strategy can be extended to other forms of inflammatory or degenerative arthritis remains unknown. Second, the present study was performed primarily in murine systems, and the translational relevance of FAP- and NCL-directed CAR-M2 cells requires validation using human FLS and human macrophages, particularly given potential differences in antigen abundance and FLS heterogeneity between mice and humans. Third, although no overt systemic or major-organ toxicity was detected, a more comprehensive safety assessment will be required to evaluate potential off-target interactions, persistence, and responses to FAP or NCL expression in non-pathological tissues. Fourth, the optimal dose, treatment frequency, route of administration, cellular persistence, and therapeutic window remain to be established. Finally, future studies should determine whether CAR-M2 therapy can provide durable disease control after a single or limited number of administrations and whether repeated treatment or combination with established anti-inflammatory or disease-modifying therapies can further enhance efficacy.

In summary, this study establishes an FLS-targeted CAR-M2 platform that integrates antigen-dependent recognition and phagocytic clearance of pathogenic FLS with IL-4-mediated macrophage immunoregulation and tissue-reparative activity. The therapeutic efficacy of both FAP- and NCL-targeted CAR-M2 cells across RA and OA models, together with the attenuation of these effects by target blockade, supports pathogenic FLS as a therapeutically actionable cellular driver shared by inflammatory and degenerative arthritis. More broadly, the reproducibility of the approach across two distinct FLS-associated targets highlights the modularity of CAR-M2 engineering and provides a framework for developing additional target-specific macrophage therapies. With further optimization of antigen selection, payload design, dosing, persistence, and safety, FLS-targeted CAR-M2 therapy may represent a promising cell-based strategy for achieving simultaneous control of synovial inflammation and progressive joint destruction in arthritis.

## Materials and Methods

### Cell culture

293FT cells (RRID: CVCL_6911) were cultured in Dulbecco’s modified Eagle’s medium (DMEM; Corning) supplemented with 10% fetal bovine serum (FBS; Corning) and penicillin-streptomycin (Sangon) at 37 °C in a humidified incubator with 5% CO₂. BMDMs were generated from bone marrow cells and used as effector cells for CAR-M2 construction^66^. Mouse FLS were maintained in RPMI 1640 medium (Corning) supplemented with 10% FBS and penicillin-streptomycin and incubated with 10 ng/ml TNFα to induce a pathogenic state^67^. For *in vitro* assays, BMDMs were seeded directly into multi-well plates and transduced with lentiviral vectors encoding CAR-M2-FAP, CAR-M2-NCL, or the corresponding control vector. Untransduced BMDMs served as BMDM-UTD controls, and BMDMs transduced with the control vector served as BMDM-vector controls. CAR expression was assessed seven days after transduction by FACS, and the resulting cells were used for co-culture, phagocytosis, RT-qPCR, RNA-seq, and downstream functional assays as indicated.

### Animals

Male C57BL/6J and DBA/1J mice (6-8 weeks old, 20-25 g) were purchased from GemPharmatech Co., Ltd., and were used for the establishment of CIA and DMM models. All the mice were housed in the Laboratory Animal Center at Southern University of Science and Technology, in a specific pathogen-free (SPF) mouse facility (room temperature, 20-22 °C; room humidity, 40-60%) with food/water access ad libitum under a 12 h light/dark cycle. All animal experiments were approved by the Institutional Animal Care and Use Committee (IACUC) of the Southern University of Science and Technology (SUSTech-JY202201007).

### Plasmid construction and virus

The FAP- and NCL-targeting CAR constructs were cloned into the pCDH-SFFV-copGFP-EF1α lentiviral vector. Each CAR construct contained an N-terminal CD8 SP, a target-specific scFv recognizing either FAP or NCL, a G4S linker, a CD8 TM, a CD3ζ intracellular signaling domain, and a Myc tag for CAR detection. GFP was co-expressed from the vector to enable tracking, enrichment, and flow cytometric analysis of transduced cells. To generate IL-4-armored CAR-M2 constructs, a T2A-murine *Il4* cassette was inserted downstream of the CAR sequence, producing a single lentiviral vector encoding the CAR and secreted IL-4. The control vector lacked the target-specific scFv and *Il4* cassette and was used to generate BMDM-vector controls. Lentiviral particles were produced by transfecting 293FT cells with the CAR or control transfer plasmid together with the packaging plasmids VPX-psPAX2 and VPX-pMD2.G using Lipofectamine 2000 (Invitrogen). The plasmids were transfected at a mass ratio of psPAX2-Vpx, pMD2.G-Vpx, and pcDNA = 3:1:4^68^. Viral supernatants were collected at 24, 48 and 72 h after transfection, filtered through a 0.45 μm membrane, and concentrated using Amicon Ultra-15 centrifugal filters (Millipore Sigma). Concentrated viral stocks were aliquoted and stored at −80 °C. Viral titers were determined using the Lenti-X p24 Rapid Titer Kit (Takara). For macrophage transduction, BMDMs were infected with lentivirus at a multiplicity of infection (MOI) of 10, and CAR expression was assessed 7 days after transduction.

### Western blot assay

Western blot analysis was performed to verify CAR expression in lentivirus-transduced BMDMs. Briefly, BMDM-UTD, CAR-M2-FAP, or CAR-M2-NCL cells were lysed, mixed with 4× loading buffer, and subjected to denaturing sodium dodecyl sulfate–polyacrylamide gel electrophoresis (SDS-PAGE). Proteins were transferred onto polyvinylidene difluoride (PVDF) membranes, and the membranes were blocked with blocking buffer before incubation overnight at 4 °C with primary antibodies, including mouse anti-Myc antibody (Abcam) to detect Myc-tagged CAR constructs and rabbit anti-Actin antibody (Beyotime) as a loading control. After washing with PBST, membranes were incubated with the corresponding secondary antibodies (Abclonal). Protein bands were visualized using the ChemiDoc Touch Imaging System (Bio-Rad).

### ELISA

IL-4 secretion by engineered BMDMs was quantified by ELISA. Briefly, culture supernatants were collected from BMDM-UTD, BMDM-vector, CAR-M2-FAP, and CAR-M2-NCL cells on day 7 after lentiviral transduction. IL-4 levels were measured using mouse IL-4 ELISA kits (Ruixin Biotechnology, Fujian, China) according to the manufacturer’s instructions. Absorbance was measured at 450 nm using an Epoch microplate reader (BioTek, USA), and IL-4 concentrations were calculated from the standard curve^69^.

### Flow cytometry

Flow cytometry was used to evaluate CAR expression and FLS phagocytosis by engineered BMDMs. Seven days after lentiviral transduction, adherent BMDMs were washed twice with PBS, detached using cell dissociation buffer or gentle scraping, and collected by centrifugation at 300 × g for 3.5 min. Cells were resuspended in PBS and analyzed using a BD FACSCanto SORP flow cytometer (BD Biosciences). CAR expression was assessed by staining BMDMs with an APC-conjugated anti-G4S linker antibody (1:1000, Cell Signaling Technology) according to the manufacturer’s instructions. G4S-positive cells were quantified in the G4S APC-A gate, with BMDM-UTD and BMDM-vector cells used as negative controls. Before the phagocytosis assays, BMDM-UTD cells were labeled with CellTracker Green (Thermo Fisher Scientific) according to the manufacturer’s instructions, enabling their identification in the FITC-A channel without relying on transduction-derived GFP. For phagocytosis assays, mouse FLS were labeled with CellTracker Red CMTPX (Thermo Fisher Scientific) according to the manufacturer’s instructions. Labeled FLS were washed twice with PBS, detached using cell dissociation buffer or gentle scraping at 37 °C for 5 min, centrifuged at 300 × g for 3.5 min, and resuspended in PBS. BMDMs were then co-cultured with labeled FLS at an effector-to-target (E:T) ratio of 3:1 for 12 h. After co-culture, cells were harvested and analyzed by flow cytometry. BMDMs and mouse FLS were detected in the FITC-A and APC-A channels, respectively, consistent with the gating strategy shown in the flow cytometry plots. Phagocytosis was evaluated as the proportion of BMDMs containing FLS-derived APC-A signal and quantified by the mean fluorescence intensity of the FLS-derived signal within the BMDM population^70^. Flow cytometry data were analyzed using FlowJo software (RRID: SCR_008520).

### RT-qPCR

BMDM-UTD, BMDM-vector, CAR-M2-FAP, or CAR-M2-NCL cells were co-cultured with mouse FLS as indicated. After co-culture, BMDMs were collected for RNA extraction. For lentivirus-transduced groups, GFP-positive BMDMs were enriched by FACS using a FACSAria SORP cell sorter (BD Biosciences), whereas BMDM-UTD controls were collected using the same macrophage gating strategy^70^. Total RNA was isolated using an Eastep Super Total RNA Extraction Kit (Promega), and cDNA was synthesized according to the manufacturer’s instructions. Quantitative PCR was performed using TB Green Premix (Takara) on a CFX96 Real-Time PCR System (Bio-Rad). Each reaction contained 0.4 μM primers and template cDNA. Relative gene expression was normalized to Gapdh and calculated using the 2^−ΔΔCt^ method, with primer sequences listed in Table S1.

### RNA-seq

For bulk RNA-seq, BMDM-vector and CAR-M2-FAP cells were co-cultured with mouse FLS under the same conditions used for the *in vitro* functional assays. After co-culture, transduced BMDMs were purified by FACS based on GFP positivity using a FACSAria SORP cell sorter (BD Biosciences), and total RNA was extracted using TRIzol reagent. RNA-seq libraries were prepared and sequenced according to standard procedures. Clean reads were aligned to the mouse reference genome using STAR, and gene-level counts were generated using featureCounts. Differential expression analysis between CAR-M2-FAP and BMDM-vector cells was performed using DESeq2. Genes with an absolute fold change ≥ 2 and a false discovery rate (FDR) < 0.05 were considered differentially expressed. GO analyses were performed using clusterProfiler.

### CIA mouse model

The CIA mouse model was established in DBA/1J mice following a previously described protocol. Bovine CII (Chondrex, USA) was emulsified with an equal volume of CFA. Mice were immunized subcutaneously at the base of the tail with 100 μL of emulsion containing 100 μg bovine CII in CFA on day 0^71^. Mice with established CIA were randomly assigned to receive PBS, BMDM-UTD, BMDM-vector, CAR-M2-FAP, CAR-M2-FAP pre-incubated with rFAP, CAR-M2-NCL, or CAR-M2-NCL pre-incubated with rNCL, as indicated in the corresponding experiments. Cells were resuspended in 200 μL PBS and administered by tail vein injection on day 28 at a dose of 5 × 10^6^ cells per mouse. Samples were collected on day 41 for μCT, histological, IF, serum biochemical, and safety analyses. Arthritis severity was evaluated by two independent blinded observers. Each paw was scored on a scale from 0 to 4 as follows: 0, normal; 1, redness and/or swelling in one joint; 2, redness and/or swelling in more than one joint; 3, redness and/or swelling of the entire paw; and 4, deformity and/or ankylosis. Scores from all paws were summed to obtain the arthritis score for each mouse. A paw score of 1 or higher was considered indicative of arthritis. Arthritis scores and body weight were monitored during the treatment period^71^.

### DMM mouse model

DMM surgery was performed on the right knee of C57BL/6J mice to induce OA, as previously described. Briefly, mice were anesthetized with 1.2% tribromoethanol, and the anteromedial meniscotibial ligament was transected to induce joint instability. Sham-operated mice underwent the same surgical exposure without ligament transection and served as controls^72^. For therapeutic studies, DMM mice received intra-articular injection on day 14 after surgery. BMDM-UTD, BMDM-vector, CAR-M2-FAP, CAR-M2-FAP pre-incubated with rFAP, CAR-M2-NCL, or CAR-M2-NCL pre-incubated with rNCL were resuspended in 20 μL PBS and injected into the right knee joint at a dose of 2 × 10^5^ cells per mouse. Samples were collected on day 35 for μCT, histological, IF, serum biochemical, and safety analyses. Body weight and general health status were monitored throughout the experimental period^73^.

### μCT analysis

Fixed, non-demineralized hind paws from CIA mice and knee joints from DMM mice were subjected to μCT analysis using a Bruker μCT system (SkyScan 1276). Samples were scanned with a 60 kV X-ray source and 100 μA current at a resolution of 10 μm. Scanned images were reconstructed using NRecon software, and three-dimensional models were generated using CTvox software. The same reconstruction parameters and threshold settings were applied across groups within each experiment. μCT reconstructions were used to evaluate bone erosion and joint destruction in CIA mice and osteophyte formation in DMM mice^42^.

### H&E staining

Paw tissues from CIA mice, knee joints from DMM mice, and major organs including the heart, lung, liver, kidney, and spleen were collected after treatment. Samples were fixed in 10% neutral-buffered formalin. Paw and knee joint tissues were decalcified in 10% EDTA (pH 7.4) at 4 °C for 40 days before paraffin embedding, whereas major organs were directly embedded in paraffin after fixation. Paraffin-embedded tissues were sectioned at 5 μm, mounted on glass slides, deparaffinized, rehydrated, and stained with H&E using standard protocols. After mounting with coverslips, stained sections were scanned using a NanoZoomer S60 digital slide scanner (Hamamatsu, Japan) and blindly evaluated by a pathologist. H&E-stained joint sections were used to assess synovial inflammation, synovial hyperplasia, and joint pathology, whereas H&E-stained major organ sections were used for safety evaluation^74^.

### SO&FG staining

Safranin O/Fast Green (SO&FG) staining was performed on paw sections from CIA mice and knee joint sections from DMM mice to assess cartilage integrity and proteoglycan content. Briefly, isolated paws and knees were fixed in 10% neutral-buffered formalin, decalcified in 10% EDTA (pH 7.4) at 4 °C for 40 days, embedded in paraffin, and sectioned at 5 μm. Sections were mounted on glass slides, deparaffinized, rehydrated, and stained with Weigert’s iron hematoxylin, followed by Fast Green staining and Safranin O counterstaining to visualize proteoglycan-rich cartilage. After dehydration and mounting, stained sections were scanned using a NanoZoomer S60 digital slide scanner (Hamamatsu, Japan) and blindly evaluated by a pathologist. For CIA mice, SO&FG-stained paw sections were used to evaluate cartilage loss and joint destruction. For DMM mice, SO&FG- and H&E-stained knee sections were used for histological assessment of OA severity, including OARSI score, cartilage area, osteophyte score, and synovitis score, according to established criteria^42^.

### IF staining

IF staining was performed on formalin-fixed, paraffin-embedded paw and knee joint sections. After deparaffinization and rehydration, sections were processed for immunostaining using the following primary antibodies: rabbit anti-COL2A1 (1:200, Abcam), mouse anti-CD80 (1:200, Cell Signaling Technology), mouse anti-IL-10 (1:200, Proteintech), mouse anti-IL-4 (1:200, Proteintech), rabbit anti-FAP (1:400, Cell Signaling Technology), and rabbit anti-NCL (1:400, Proteintech). After incubation with appropriate fluorescence-conjugated secondary antibodies, nuclei were counterstained with DAPI-containing mounting medium (Beyotime, China). Images were captured using a TCS SP8 confocal microscope (Leica) equipped with a 20× objective lens. For CIA paw sections and DMM knee sections, co-localization of FAP or NCL with GFP-positive CAR-M2 was quantified as the co-localization ratio. Paw sections from CIA mice were used to quantify CD80-, IL-10-, IL-4-, FAP-, and NCL-positive cells. Knee sections from DMM mice were used to quantify COL2A1-, CD80-, IL-10-, IL-4-, FAP-, and NCL-positive cells. Quantification was performed under the same imaging and threshold settings across groups^75^.

### Serum assays

At the experimental endpoint, mouse serum was collected for biochemical safety analysis. Serum levels of ALB, ALP, ALT, AST, urea, and Cr were measured using an automatic biochemical analyzer (MS-480, Medical System). These parameters were used to assess potential liver and kidney toxicity after CAR-M2 treatment^70^.

### Statistical analysis

GraphPad Prism software (RRID: SCR_002798) was used for graph generation and statistical analyses. All *in vitro* experiments were performed at least three times independently using BMDM batches generated from different mouse donors, unless otherwise specified. For *in vivo* experiments, n = 5 mice per group. Data are presented as mean ± SD. For comparisons among multiple groups, one-way ANOVA followed by Tukey’s post hoc test was used for all pairwise comparisons. For comparisons between two matched groups, a two-tailed paired Student’s t test was used. Time-course arthritis scores were analyzed using two-way repeated-measures ANOVA followed by Tukey’s post hoc test. Sample size was determined based on previous studies and pilot experiments. *P* < 0.05 was considered statistically significant.

## Acknowledgements

The authors acknowledge the assistance of the Core Research Facilities, Microscope and Imaging Center, and Experimental Animal Center of the Southern University of Science and Technology. The authors also acknowledge the use of ChatGPT for language refinement and spell-checking to improve manuscript readability and take full responsibility for the final content.

## Contributors

C.L., X.Y., and L.Z. conceived the study. X.Y., C.L., L.Z., and Y.X. designed the experiments. Y.X. and Z.W. performed experiments. D.X. assisted with the experiment. Z.W. and Y.X. wrote the manuscript. C.L., X.Y., and L.Z. revised the manuscript. C.L., X.Y., and L.Z. supervised and funded the study.

## Data Sharing Statement

The data that support the findings of this study are available from the corresponding author upon reasonable request.

## Supplementary Information

**Figure S1.**
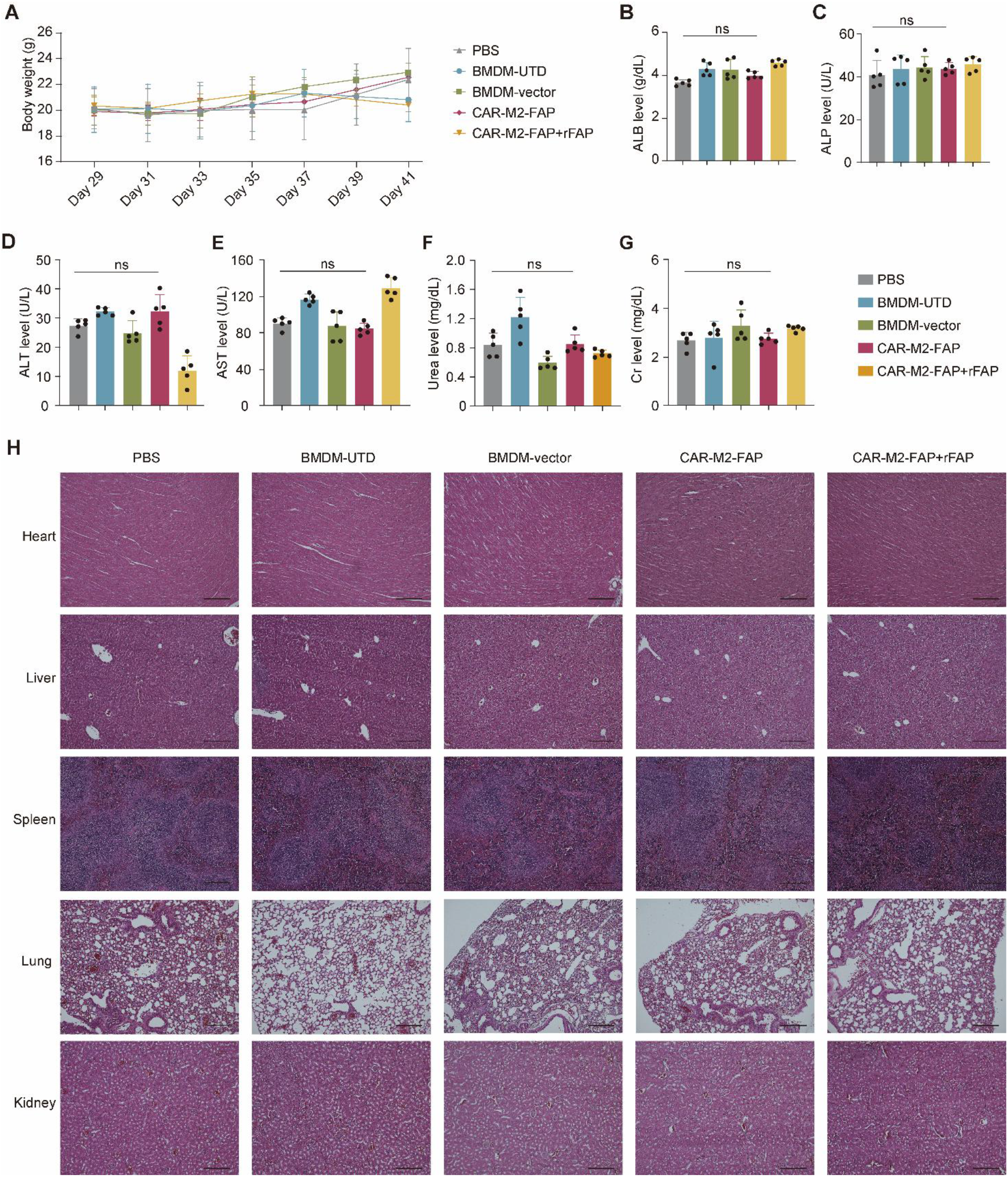
Safety evaluation of CAR-M2-FAP in CIA mice. **(A)** Changes in body weight of CIA mice following the indicated treatments. **(B-G)** Serum levels of ALB **(B)**, ALP **(C)**, ALT **(D)**, AST **(E)**, urea **(F)**, and Cr **(G)**, as measured using an automated biochemical analyzer. **(H)** Representative H&E-stained sections of major organs from the indicated treatment groups. Scale bar = 300 μm. Data are presented as mean ± SD. n = 5 per group. *P-*values were calculated by one-way ANOVA followed by Tukey’s post hoc test **(B-G)**. ns, not significant.

**Figure S2.**
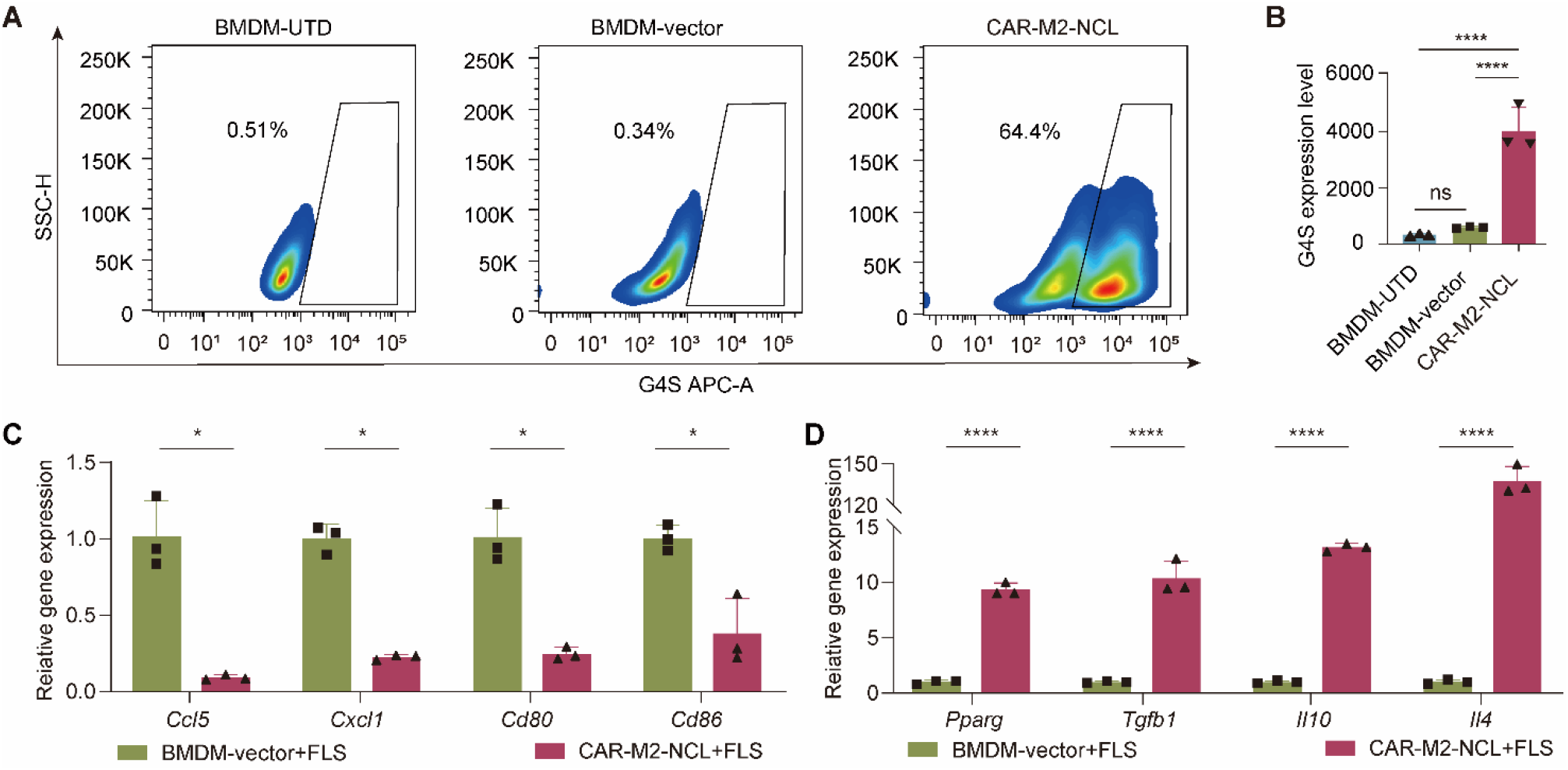
*In vitro* characterization of CAR-M2-NCL. **(A** and **B)** Representative flow cytometry plots **(A)** and quantification **(B)** of G4S expression in BMDMs. **(C** and **D)** RT-qPCR analysis of M1-associated and pro-inflammatory genes **(C)** and M2-associated and anti-inflammatory genes **(D)** in BMDMs after co-culture with FLS. Each experiment was repeated three times. Data are presented as mean ± SD. *P-*values were calculated by one-way ANOVA followed by Tukey’s post hoc test **(B)** or paired Student’s t-test **(C** and **D)**. \**P* < 0.05, \*\*\*\**P* < 0.0001, ns, not significant.

**Figure S3.**
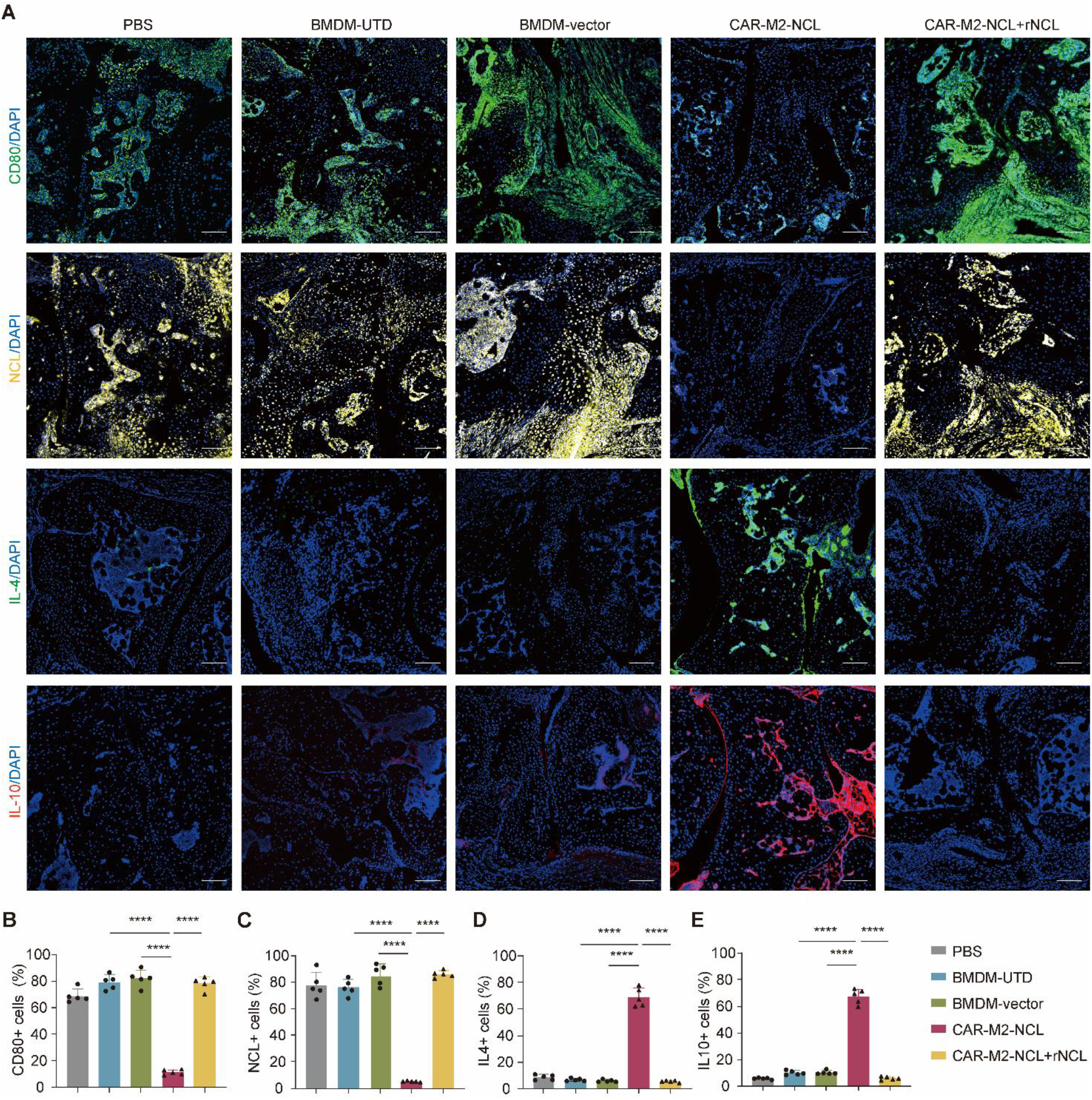
IF analysis of CAR-M2-NCL-treated CIA mice. **(A)** Representative IF staining of paw sections from different treatment groups using an anti-CD80, anti-NCL, anti-IL-4, or anti-IL-10 antibody. Nuclei were counterstained with DAPI. Scale bar = 50 μm. **(B-E)** Quantification of CD80-positive cells **(B)**, NCL-positive cells **(C)**, IL-4-positive cells **(D),** and IL-10-positive cells **(E)** in paw sections from different treatment groups. Data are presented as mean ± SD. n = 5 per group. *P*-values were calculated by one-way ANOVA followed by Tukey’s post hoc test **(B-E)**: \*\*\*\**P* < 0.0001, ns, not significant.

**Figure S4.**
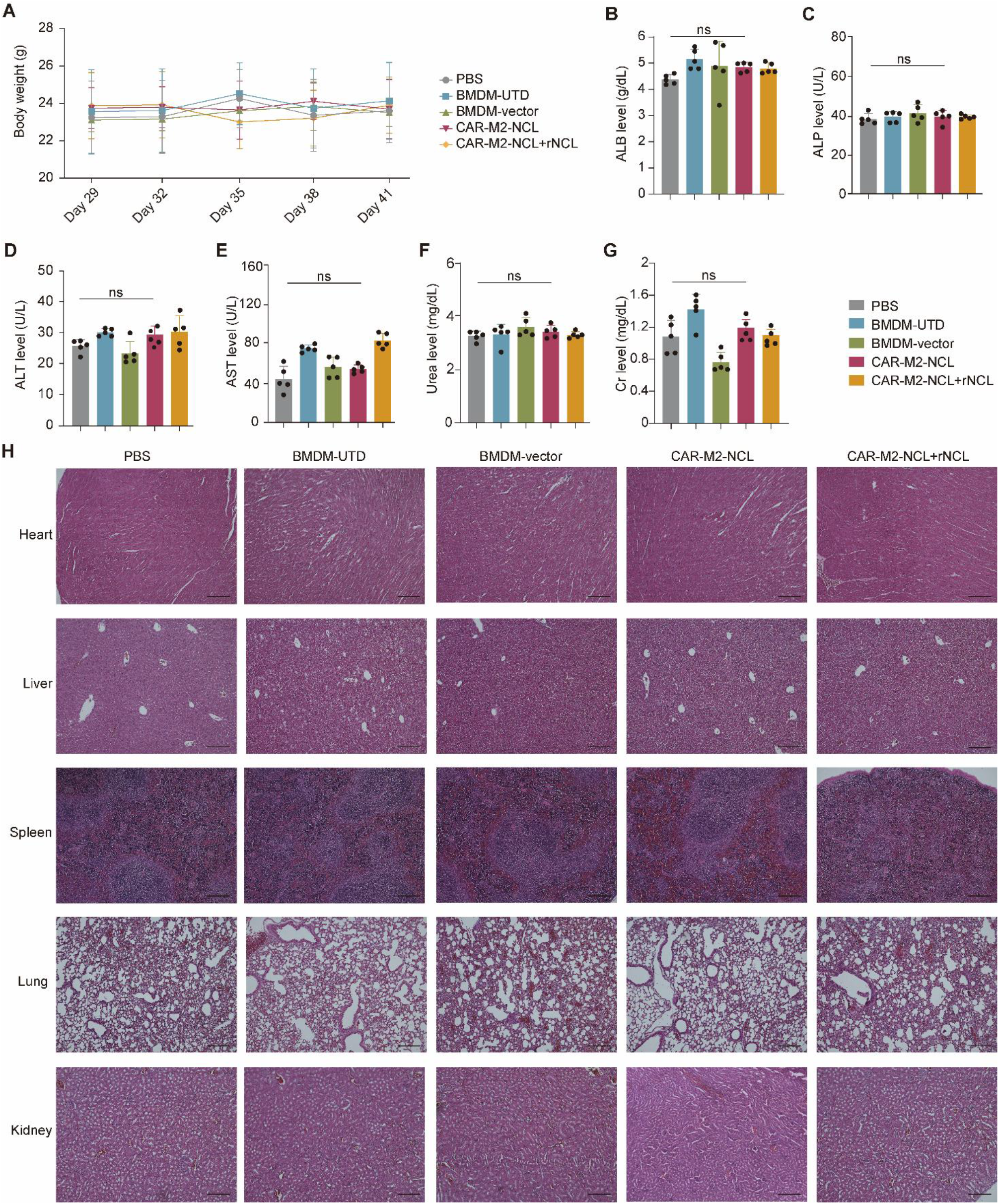
Safety evaluation of CAR-M2-NCL in CIA mice. **(A)** Body weight changes in CIA mice after different treatments. **(B-G)** Serum levels of ALB **(B)**, ALP **(C)**, ALT **(D)**, AST **(E)**, urea **(F),** and Cr **(G)**, as determined by an automated biochemical analyzer. **(H)** H&E staining of major organs from different treatment groups. Scale bar = 300 μm. Data are presented as mean ± SD. n = 5 per group. *P-*values were calculated by one-way ANOVA followed by Tukey’s post hoc test **(B-G)**. ns, not significant.

**Figure S5.**
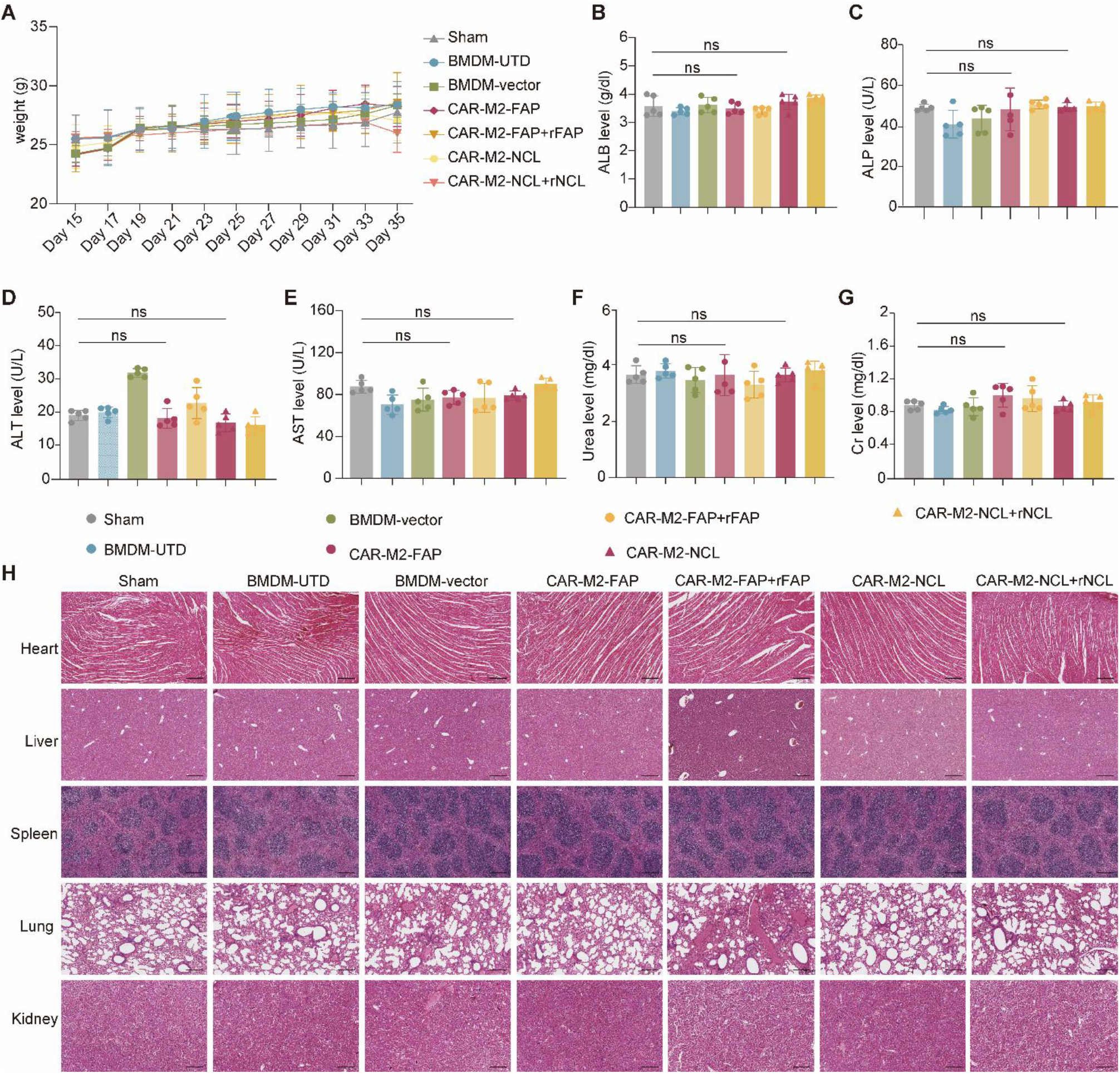
Safety evaluation of CAR-M2 in DMM mice. **(A)** Body weight changes in DMM mice after different treatments. **(B-G)** Serum levels of ALB **(B)**, ALP **(C)**, ALT **(D)**, AST **(E)**, urea **(F)**, and Cr **(G)**, as determined by an automated biochemical analyzer. **(H)** H&E staining of major organs from different treatment groups. Scale bar = 300 μm. Data are presented as mean ± SD. n = 5 per group. *P-*values were calculated by one-way ANOVA followed by Tukey’s post hoc test **(B-G)**. ns, not significant.

**Table S1.**
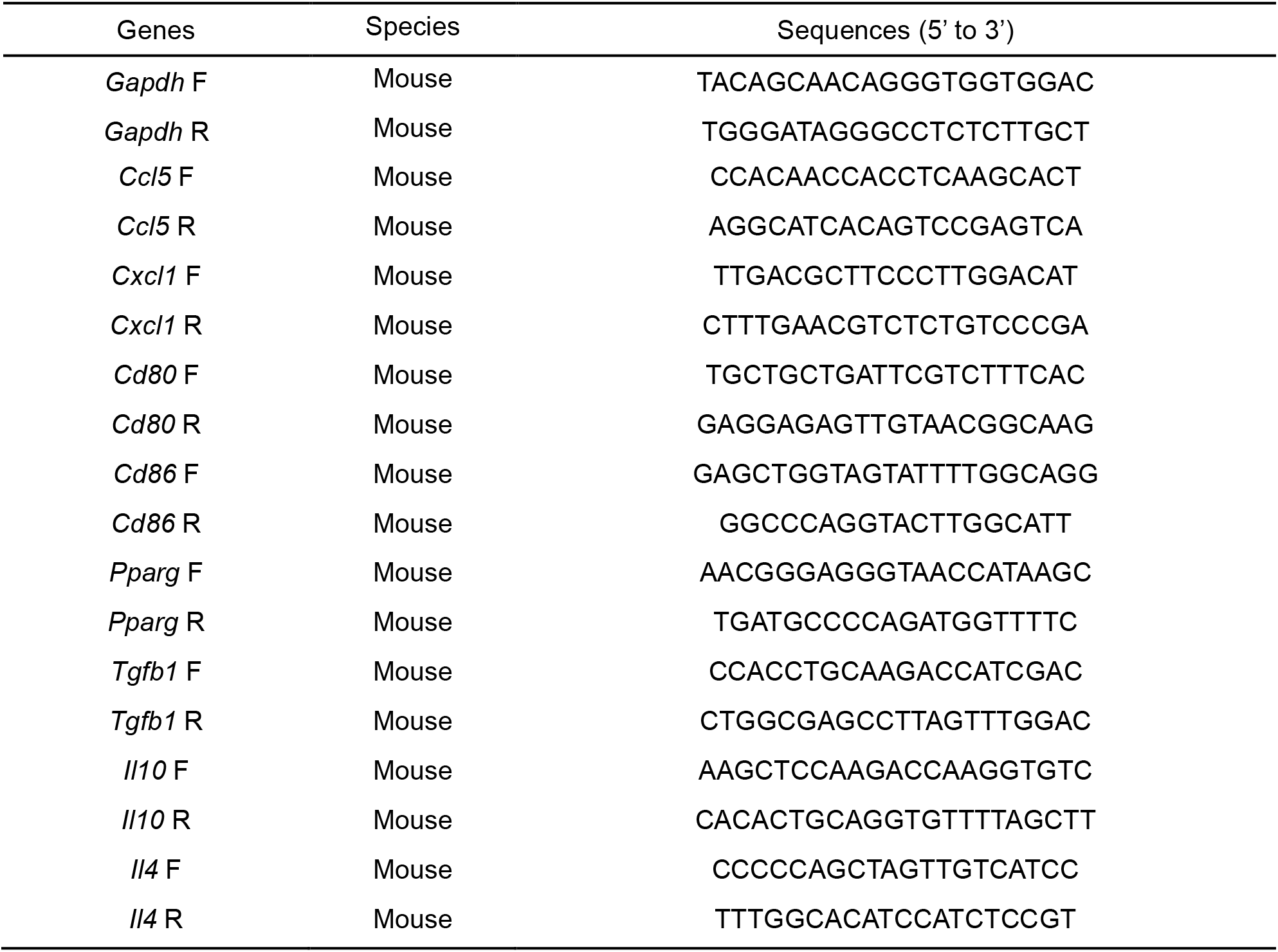
Primer sequence list for RT-qPCR.

