## Supplementary Information for "CAR-macrophage immunotherapy targeting fibroblast-like synoviocytes ameliorates inflammatory and degenerative arthritis"

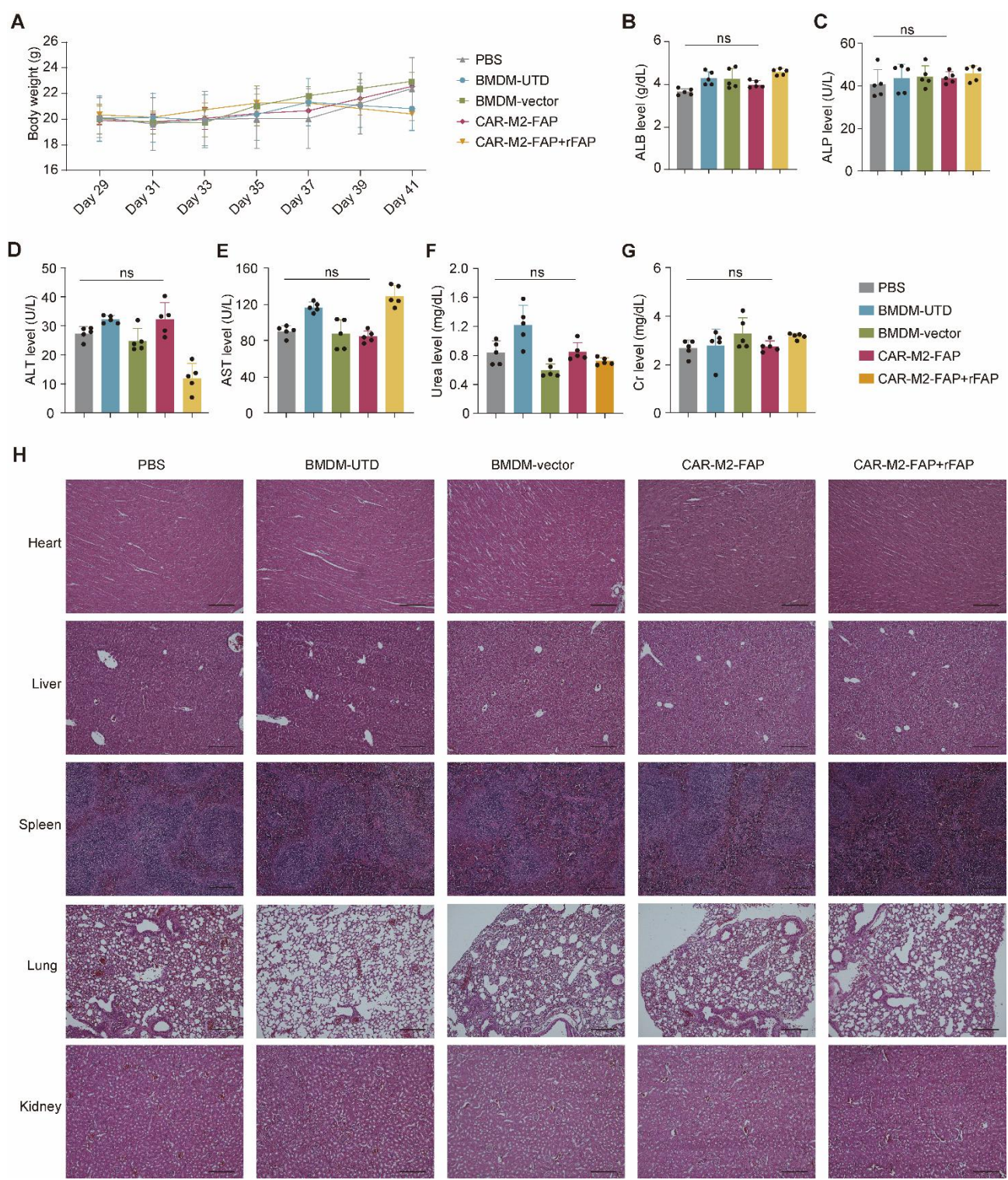

**Figure S1. Safety evaluation of CAR-M2-FAP in CIA mice.** (A) Changes in body weight of CIA mice following the indicated treatments. (B-G) Serum levels of ALB (B), ALP (C), ALT (D), AST (E), urea (F), and Cr (G), as measured using an automated biochemical analyzer. (H) Representative H&E-stained sections of major organs from the indicated treatment groups. Scale bar = 300 μm. Data are presented as mean ± SD. $n = 5$ per group. $P$-values were calculated by one-way ANOVA followed by Tukey's post hoc test (B-G). ns, not significant.

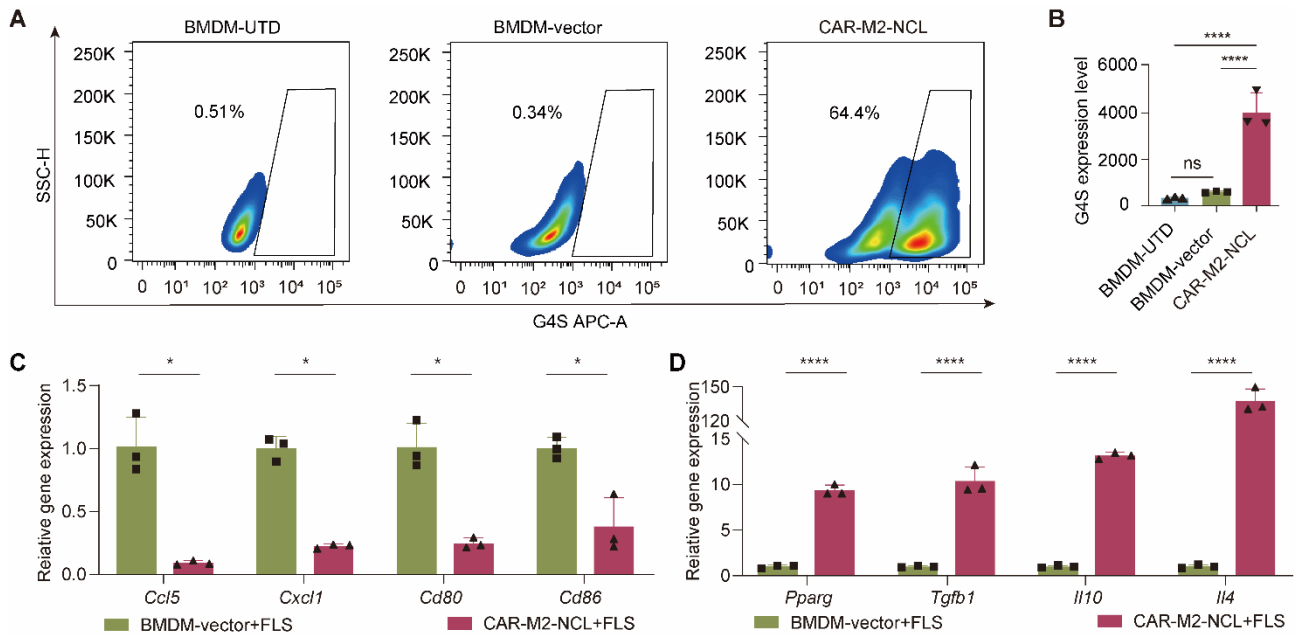

**Figure S2. *In vitro* characterization of CAR-M2-NCL.** (A and B) Representative flow cytometry plots (A) and quantification (B) of G4S expression in BMDMs. (C and D) RT-qPCR analysis of M1-associated and pro-inflammatory genes (C) and M2-associated and anti-inflammatory genes (D) in BMDMs after co-culture with FLS. Each experiment was repeated three times. Data are presented as mean ± SD. *P*-values were calculated by one-way ANOVA followed by Tukey's post hoc test (B) or paired Student's t-test (C and D). \**P* < 0.05, \*\*\*\**P* < 0.0001, ns, not significant.

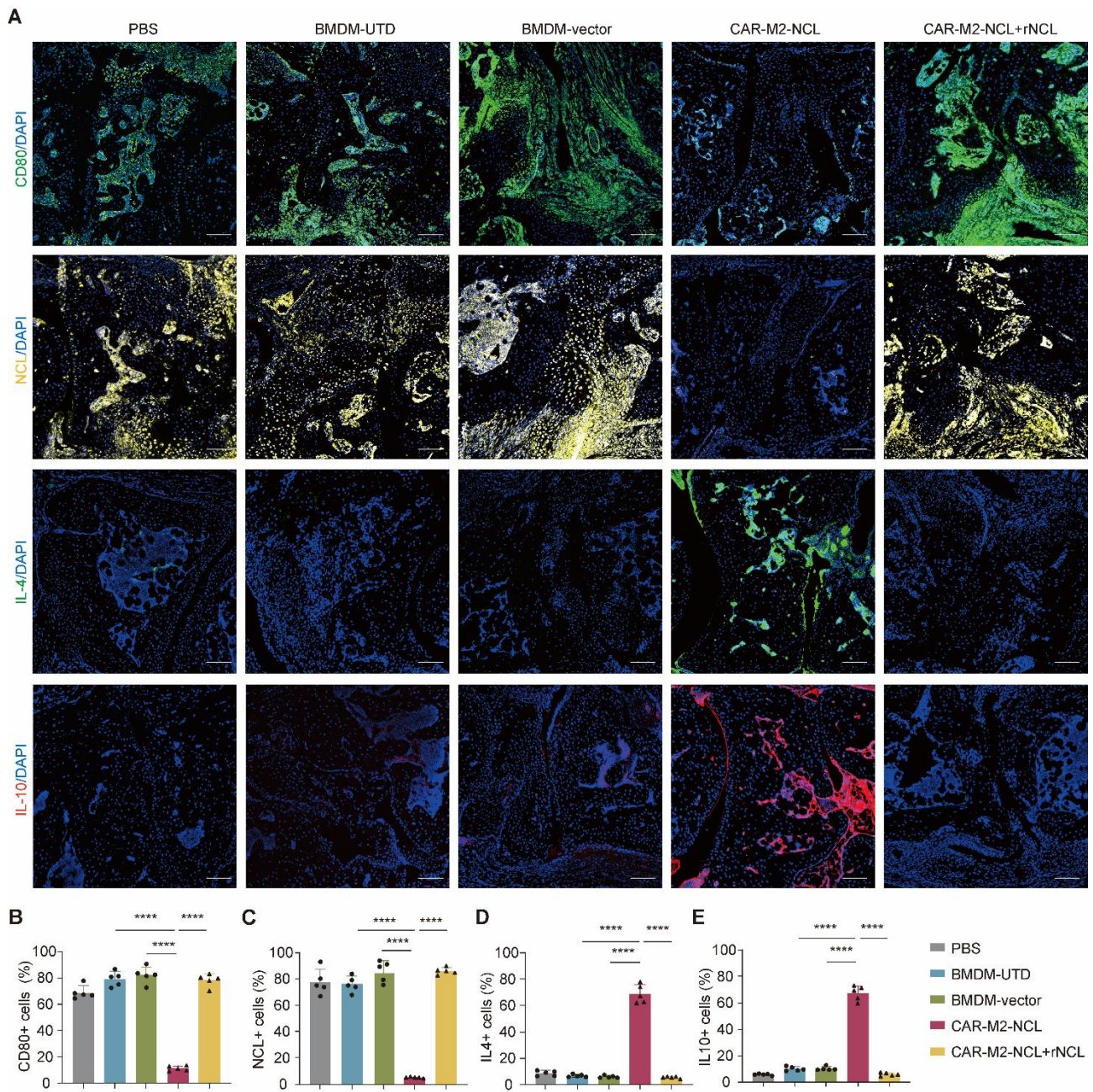

**Figure S3. IF analysis of CAR-M2-NCL-treated CIA mice. (A)** Representative IF staining of paw sections from different treatment groups using an anti-CD80, anti-NCL, anti-IL-4, or anti-IL-10 antibody. Nuclei were counterstained with DAPI. Scale bar = 50 μm. **(B-E)** Quantification of CD80-positive cells **(B)**, NCL-positive cells **(C)**, IL-4-positive cells **(D)**, and IL-10-positive cells **(E)** in paw sections from different treatment groups. Data are presented as mean ± SD. $n = 5$ per group. $P$-values were calculated by one-way ANOVA followed by Tukey's post hoc test **(B-E)**: \*\*\*\*$P < 0.0001$, ns, not significant.

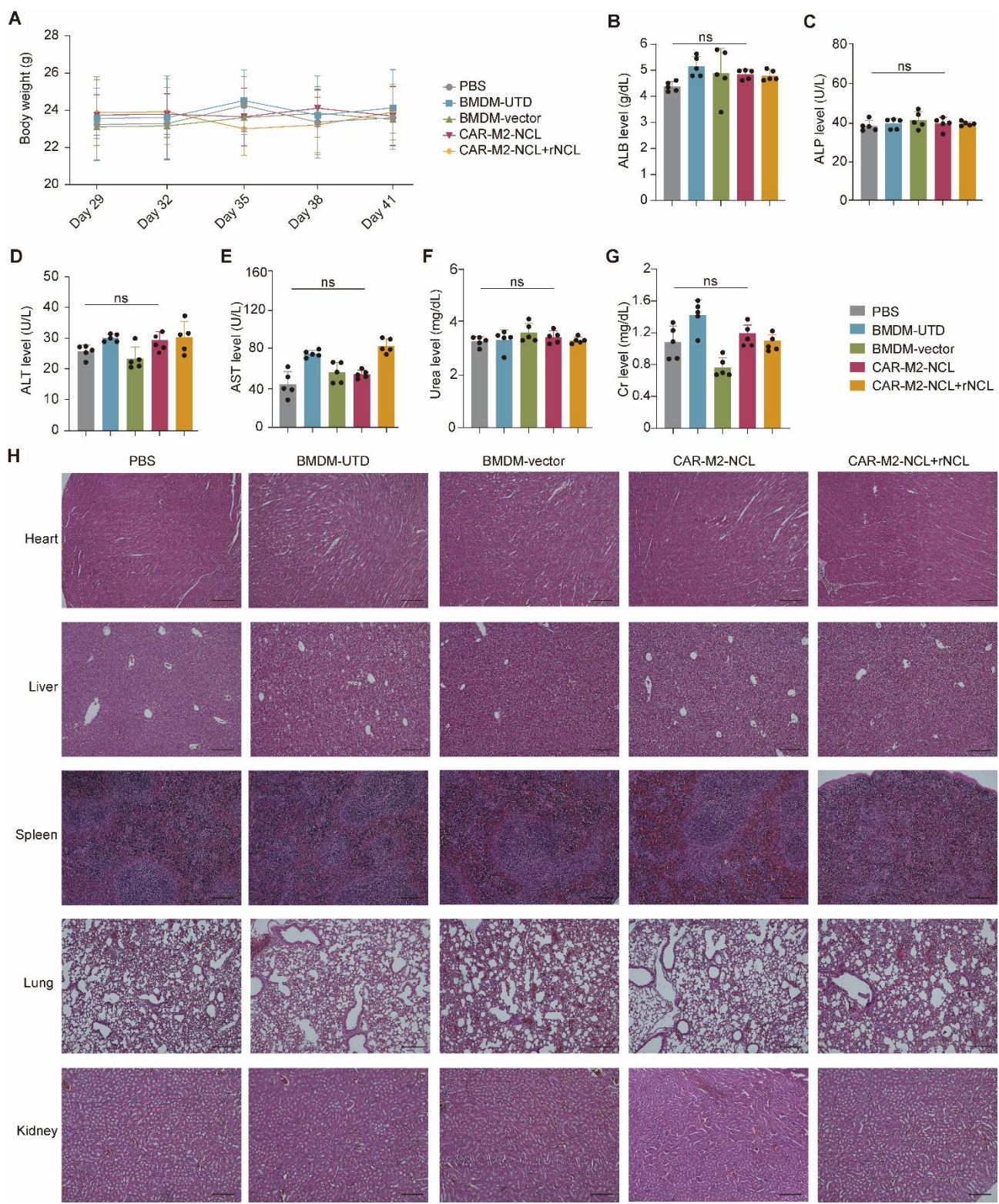

**Figure S4. Safety evaluation of CAR-M2-NCL in CIA mice.** (A) Body weight changes in CIA mice after different treatments. (B-G) Serum levels of ALB (B), ALP (C), ALT (D), AST (E), urea (F), and Cr (G), as determined by an automated biochemical analyzer. (H) H&E staining of major organs from different treatment groups. Scale bar = 300 μm. Data are presented as mean ± SD. $n = 5$ per group. $P$-values were calculated by one-way ANOVA followed by Tukey's post hoc test (B-G). ns, not significant.

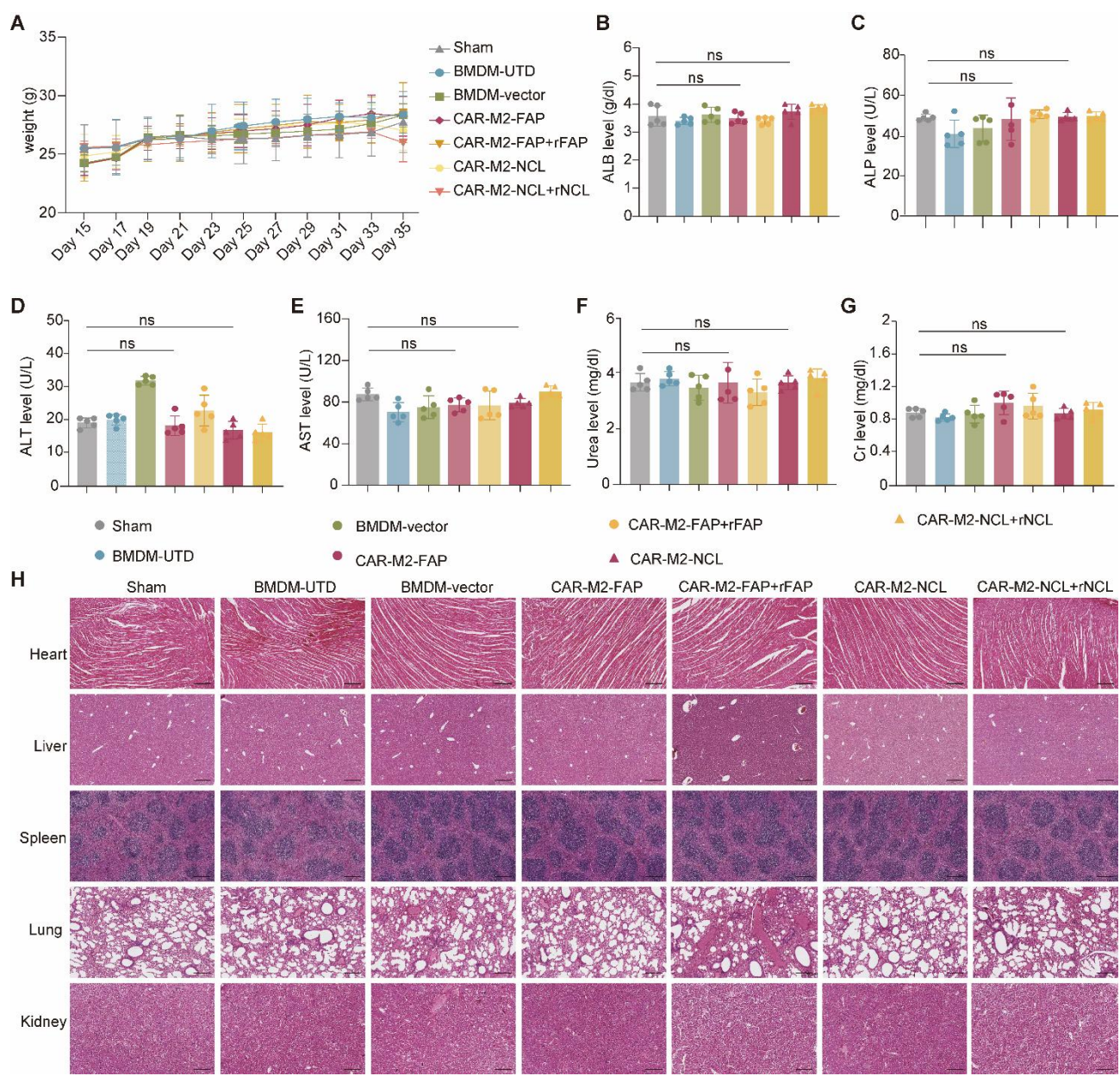

**Figure S5. Safety evaluation of CAR-M2 in DMM mice.** (A) Body weight changes in DMM mice after different treatments. (B-G) Serum levels of ALB (B), ALP (C), ALT (D), AST (E), urea (F), and Cr (G), as determined by an automated biochemical analyzer. (H) H&E staining of major organs from different treatment groups. Scale bar = 300 μm. Data are presented as mean ± SD. $n = 5$ per group. $P$-values were calculated by one-way ANOVA followed by Tukey's post hoc test (B-G). ns, not significant.

54 **Table S1. Primer sequence list for RT-qPCR**

| Genes | Species | Sequences (5' to 3') |
| --- | --- | --- |
| *Gapdh* F | Mouse | TACAGCAACAGGGTGGTGGAC |
| *Gapdh* R | Mouse | TGGGATAGGGCCTCTCTTGCT |
| *Ccl5* F | Mouse | CCACAACCACCTCAAGCACT |
| *Ccl5* R | Mouse | AGGCATCACAGTCCGAGTCA |
| *Cxcl1* F | Mouse | TTGACGCTTCCCTTGGACAT |
| *Cxcl1* R | Mouse | CTTTGAACGTCTCTGTCCCGA |
| *Cd80* F | Mouse | TGCTGCTGATTCGTCTTTCAC |
| *Cd80* R | Mouse | GAGGAGAGTTGTAACGGCAAG |
| *Cd86* F | Mouse | GAGCTGGTAGTATTTTGGCAGG |
| *Cd86* R | Mouse | GGCCCAGGTACTTGGCATT |
| *Pparg* F | Mouse | AACGGGAGGGTAACCATAAGC |
| *Pparg* R | Mouse | TGATGCCCCAGATGGTTTTC |
| *Tgfb1* F | Mouse | CCACCTGCAAGACCATCGAC |
| *Tgfb1* R | Mouse | CTGGCGAGCCTTAGTTTGGAC |
| *Il10* F | Mouse | AAGCTCCAAGACCAAGGTGTC |
| *Il10* R | Mouse | CACACTGCAGGTGTTTTAGCTT |
| *Il4* F | Mouse | CCCCCAGCTAGTTGTCATCC |
| *Il4* R | Mouse | TTTGGCACATCCATCTCCGT |
